# Innate Immune Memory is Stimulus Specific

**DOI:** 10.1101/2025.01.22.634275

**Authors:** Aoife O’Farrell, Zijian Niu, Jingxin Li, Laura C. Van Eyndhoven, Kavitha Sarma, Arjun Raj

## Abstract

Innate immune memory (also termed trained immunity) is defined in part by its ability to cross-protect against heterologous pathogens, and can be generated by many different stimuli, suggesting a “universal” trained state. However, different stimuli could form distinct memories, leading to stimulus-specific trained responses. Here, we use primary human monocyte-derived macrophages to demonstrate phenotypic and epigenetic stimulus specificity of innate immune memory six days after initial exposure. Quantification of cytokine production with single-molecule RNA imaging demonstrates stimulus-specific patterns of response to restimulation at the single cell level. Differential licensing of inflammatory transcription factors is associated with encoding of specificities in chromatin. Trained cells show stronger responses to secondary stimuli that are more similar to the initial stimulus they experienced, suggesting a functional role for these stimulus-specific memories. Rather than activating a universal training state, our findings demonstrate that different stimuli impart specific memories that generate distinct training phenotypes in macrophages.

## INTRODUCTION

Although once believed incapable of forming memory, macrophages are now understood to change functionally following an encounter with an inflammatory stimulus in a process termed “trained immunity” or, more broadly, “innate immune memory”^1–3^. This memory is currently conceptualized as nonspecific in the sense that cells are not thought to retain information about the initial stimulus they experienced. Rather, the ability to cross-protect, whereby memory of one pathogen allows for a stronger and faster response to a heterologous pathogen, seemingly argues against specificity in the induced memory. Specificity, on the other hand, would manifest as the same cells showing different secondary responses depending on the nature of the initial stimulus. It remains unknown whether such differences exist.

Current classification criteria for innate immune memory describe largely qualitative differences in secondary responses due to a memory induced by a primary stimulus. At the broadest level, innate immune memory in macrophages can lead to a subsequent secondary response that is either stronger (trained) or weaker (tolerant) than the primary response^3–5^. While trained vs. tolerant responses are qualitatively different behaviors that can arise from different initial stimuli, providing some evidence for specificity, it has generally been assumed that within the subset of stimuli that lead to trained responses, there is not much difference in the trained response arising from different stimuli. This conclusion stems from measuring trained responses via qualitative changes to phenotype compared to untrained cells, such as functional protection from infection^6–8^, metabolic changes^9,10^, or increased production of key cytokines such as *TNF* and *IL6*^11,12^. Much effort has thus gone into identifying the stimuli capable of generating this seemingly “universal” trained state, including *in vitro* pathogen stimulus^5,13,14^, vaccination^8,11,15^, infection *in vivo*^16,17^, host factors^12,18^, and tissue injury^19,20^. Comparatively less focus, however, has been placed on whether all training stimuli generate the *same* trained state—that is, how the use of different training stimuli may lead to differences in canonical trained phenotypes.

Recent high-dimensional analyses have indeed hinted at the possibility of different training stimuli leading to different internal states. For instance, the primary (untrained) response itself can show specificity^21–23^, raising the possibility that cells might remember the particulars of a stimulus-specific immune response after it has concluded. Indeed, some prior work does suggest specificity of macrophage memory at short timescales (around 24 hours after stimulation)^24–27^ however, at that timescale, those differences may still reflect aspects of the primary response, prompting us to wonder whether such specificities could still be present after that transient initial response has dissipated, which would represent a longer-lasting stimulus-specific memory.

It is also unclear whether specificity in innate immune memory is encoded in particular subsets of cells. Observations of marked single-cell variation of gene expression within macrophages^28–31^ raises the possibility that different individual macrophages within the population may themselves have different abilities to form innate immune memories. Quantitatively assessing this possibility requires accurate and sensitive measurements of expression levels in individual cells under different primary and secondary stimuli.

Here, we used high-resolution single-cell imaging approaches and bulk epigenetic profiling to identify specificities in memory formation and maintenance by distinct training stimuli (β-Glucan, MDP, IFNγ) in primary human monocyte-derived macrophages. We quantified both the encoding and phenotype of memory of distinct stimuli and identified consistent stimulus-specific patterns in cytokine production, morphology, and functional ability of macrophages depending on their training stimulus. We measured differences in transcription factor binding in regions of accessible chromatin in cells trained with different stimuli, identifying distinct patterns of transcription factor activation and durability of chromatin accessibility changes. We used single-cell cytokine expression data to demonstrate that any single macrophage is capable of training. By measuring the trained response of cells from the same donor several times, we suggest that inter-individual variation is primarily due to transient nongenetic states. Overall, our results quantify stimulus specificity of memory in macrophages and suggest an element of “learning” resulting from stimulus-specific memories of prior experiences.

## RESULTS

### There is not a discrete universal trained state

We wanted to determine whether different training stimuli elicited the same training state. As such, we first looked for differences in response to secondary stimulation between cells trained with distinct stimuli. We elected to use *in vitro* human monocyte-derived macrophages, a commonly used model system for innate immune memory^5,13,17,31,32^. We chose to examine two different primary stimuli in detail: the fungal ligand β-Glucan, which is well established to generate trained immunity *in vitro*^4,5,33^; and the bacterial mimetic muramyl dipeptide (MDP), which is thought to mimic the cellular pathways induced through *in vivo* training with the BCG vaccine^15,34^. β-Glucan and MDP are both commonly used to induce innate immune memory but signal through distinct pathways–as such, comparing the trained responses they generate allowed us to test the hypothesis that memories specific to the primary stimulus can affect the secondary response.

We collected monocytes from the apheresis product of healthy donors and stimulated them with primary stimulus (β-Glucan or MDP) for 24 hours. Untrained (control) cells received no stimulation during this time. After 24 hours, β-Glucan or MDP was removed, and the cells were allowed to rest for five additional days. On day six, we evaluated the phenotype of these cells, both before and after a secondary stimulation as a readout of cell activity (**Figure 1A**). Before secondary stimulation on day 6, trained cells were no longer producing proinflammatory cytokines (**Figure 1B, Supplemental Figure 1A**); this reversion to baseline levels is archetypal of the trained phenotype^3^.

**Figure 1:**
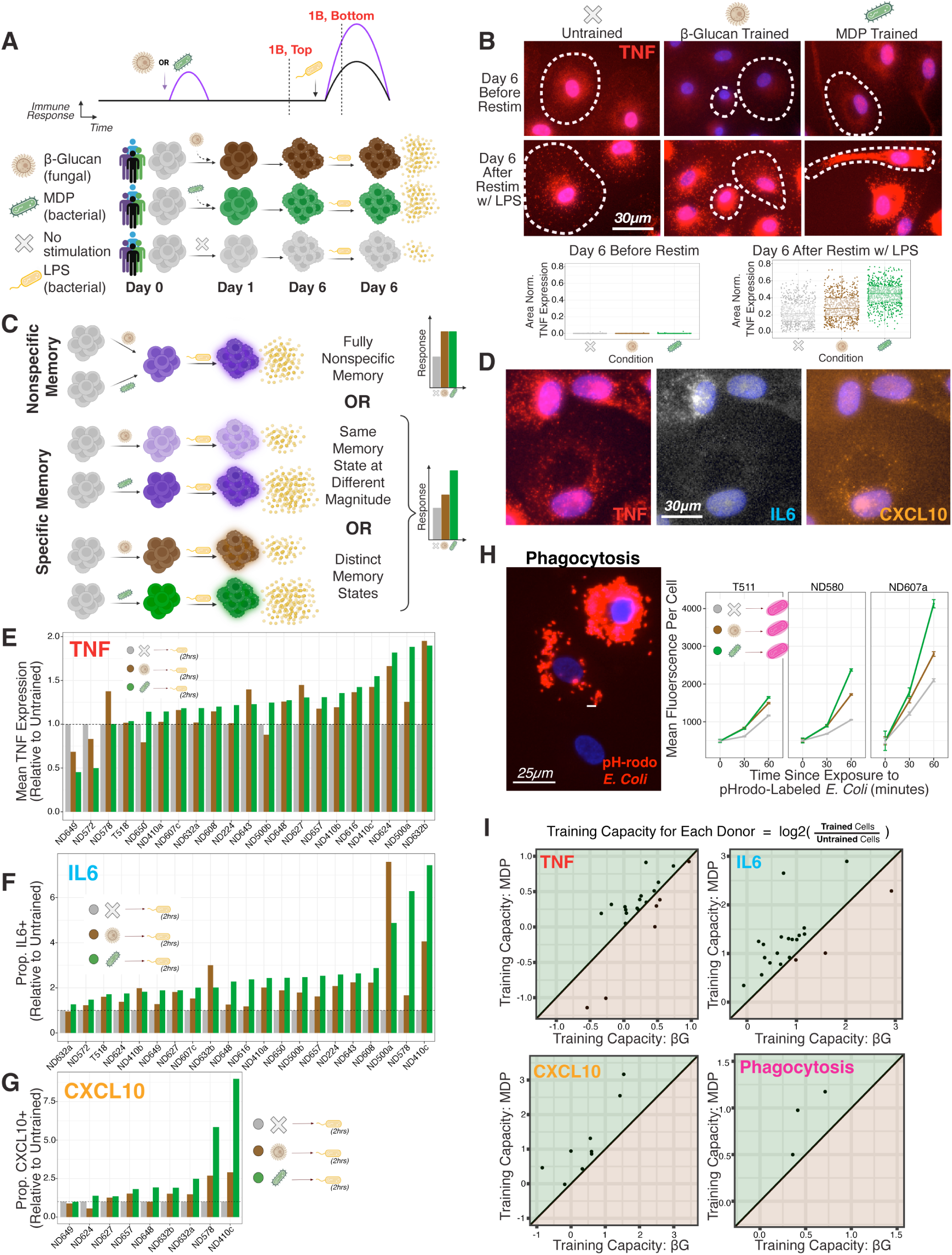
There is Not a Discrete Universal Trained State. **(A)** Schematic of experimental procedure. Human monocytes were stimulated with either fungal protein β-Glucan or bacterial mimetic MDP on the first day of culture, before resting for five days. The cytokine response to secondary stimulation with heterologous pathogen LPS was measured by single-molecule RNA FISH. **(B) *Top***: Representative single-molecule RNA FISH images of *TNF* expression before (top) and two hours after (bottom) stimulation with 100ng/mL lipopolysaccharide (LPS). Each individual spot is one *TNF* transcript. ***Bottom:*** Quantified data from experiment shown above, showing TNF expression per cell for untrained (grey), β-Glucan-trained (brown), and MDP-trained (green) populations. Each dot shows the *TNF* transcripts per square micron for an individual cell. Donor shown is ND500a. **(C)** Schematic of possible results for quantified gene expression. If the response to secondary stimulation for both train conditions matches each other, memory is fully nonspecific. If responses for the two train conditions differ from each other, memory is stimulus-specific. **(D)** Representative single-molecule RNA FISH images of *TNF, IL6,* and *CXCL10* expression two hours after stimulation with 100ng/mL LPS. Donor shown is ND648. **(E)** Mean area-normalized *TNF* transcripts per cell in untrained (grey), β-Glucan-trained (brown), or MDP-trained (green) populations, two hours after stimulation with LPS. Values are normalized to the untrained value for that donor (indicated by dashed line). Approximately 400 single cells were analyzed per condition **(F-G)** Proportion of first responder cells expressing *IL6* or *CXCL10* in untrained (grey), β-Glucan-trained (brown), or MDP-trained (green) populations, two hours after stimulation with LPS. Values are normalized to the untrained value for that donor (indicated by dashed line). Approximately 400 single cells were analyzed per condition. **(H) *Left:*** representative image of fluorescent signal from cells 1 hour after stimulation with pH-rodo labeled *E. coli*. Donor shown is ND580. ***Right:*** Mean fluorescence per cell (as a metric for amount of phagocytosed particles) over time for untrained (grey), β-Glucan-trained (brown), or MDP-trained (green) cells. Error bars show standard error of the mean for approximately 350 single cells analyzed per condition. **(I)** Summary plots of results from E-H, showing training capacity for each donor in β-Glucan versus MDP trained cells. Training capacity was calculated by the log2 fold change of the difference in mean area-normalized transcripts per cell (*TNF*), the difference in proportion of first responders (*IL6, CXCL10*), or the difference in mean fluorescence per cell (phagocytosis) between trained and untrained cells. Diagonal *y=x* line denotes where “nonspecific training” would lie on these plots, with points above the line showing stronger training from MDP, and points below showing stronger training from β-Glucan.

Differences in the response to secondary stimulation between cells that received a primary stimulation (trained cells) six days prior vs. the control cells that did not receive primary stimulus (untrained cells) indicates that the cells have committed at least some aspect of the initial response to cellular memory, forming the basis of the trained response. This memory is non-specific if we cannot distinguish the primary stimulus type by evaluating the memory state thereafter; that is, if cells that experienced different primary stimuli appear identical after some period of time and also respond identically upon secondary stimulation. However, if there are differences in the memory state as a function of the primary stimulus, there is by definition some element of specificity of the memory. Specificity could come in the form of differences in magnitude of a general state (with some stimuli forming a “stronger” memory further distinct from untrained cells) or in differences in the state itself (with different stimuli encoding distinct memories) **(Figure 1C)**.

We began our search for training specificity by looking for differences in cytokine and chemokine expression (*TNF, IL6, CXCL10*) in trained and untrained cells two hours after secondary stimulation with bacterial lipopolysaccharide (LPS), a potent inflammatory activator that is recognized by TLR4, which does not respond to either of our trained stimuli. We used single-molecule RNA fluorescence *in situ* hybridization (single-molecule RNA FISH) to compare expression within the same donor after secondary stimulation, allowing for highly accurate quantification of cytokine/chemokine transcription in absolute molecular units with single-cell resolution. This method provided quantitative assessment of expression differences between trained and untrained cells. Two hours after stimulation with LPS, almost every macrophage produced *TNF*, while only a subset of “first responders” produced *IL6* (∼10% of untrained cells) or *CXCL10* (∼40% of untrained cells) (**Figure 1D**). Given the relatively uniform induction of *TNF* transcription, we quantified expression of that gene by counting the number of transcripts per cell (normalized by cell area^35^). For *IL6* and *CXCL10*, only a small proportion of cells showed signs of active transcription, so for those genes we elected to measure the proportion of responding cells in each population that were expressing these genes; however, we also observed training for *IL6* and *CXCL10* when measuring area-normalized transcripts per cell (**Supplemental Figure 1E**).

All primary stimuli showed evidence of training for all measured genes in the majority of donors, meaning that expression was higher in trained cells than untrained cells two hours after stimulation with LPS (**Figure 1E-G, Supplemental Figure 1C-D**). Altogether, 20 of 21 of donors showed significantly higher cytokine levels in at least one training condition compared to untrained cells for *IL6* expression (8 of 9 donors for *CXCL10*, and 18 of 21 for *TNF*). Increased local cell density was not associated with the proportion of cells expressing *IL6* or *CXCL10* in trained or untrained populations, suggesting cell-cell communication did not affect the number of first responders at this early time point (**Supplemental Figure 2A**).

We next looked for specificity by measuring the differences in the amount of training produced by different primary stimuli. For the three genes we measured, MDP-trained cells consistently expressed more cytokine and chemokine than β-Glucan-trained cells after stimulation with LPS (**Figure 1E-G**). In 15 of 21 donors (71%), cells trained with β-Glucan or MDP had significantly different *IL6* production from each other, with MDP-trained cells showing significantly higher expression than β-Glucan-trained cells in 62% of total donors. We saw similar trends in *CXCL10* (8 of 9 donors with differential expression between β-Glucan- and MDP-trained cells, 89% of all donors MDP highest) and *TNF* expression (16 of 21 donors with differential expression between β-Glucan- and MDP-trained cells, 52% of all donors MDP highest). Across three genes and 21 donors, we saw a consistent stimulus-specific pattern to memory responses, in which cells trained with MDP expressed more cytokine and chemokine than cells trained with β-Glucan when restimulated with LPS.

We wondered whether specific responses extended beyond the first responder window to a longer timescale following stimulation with a heterologous stimulus. We performed a timecourse analysis of *TNF* and *IL6* expression in trained and untrained cells after secondary stimulation with LPS **(Supplemental Figure 2B-C)**. Both β-Glucan- and MDP-trained cells responded faster than untrained cells initially, but expression of *TNF* and *IL6* in β-Glucan-trained cells eventually dropped below that of untrained cells. Expression of *TNF* and *IL6* in MDP-trained cells remained greater than untrained cells in one donor, and matched the level of untrained cells in one donor. These results show that trained responses to secondary stimulation have stimulus-specific trajectories, and suggest that training may lead to faster, but not necessarily stronger, transcription of inflammatory cytokines and chemokines.

Macrophages perform many tasks beyond just cytokine and chemokine production. One of these is phagocytosis, the ability to internalize pathogens. We measured the ability of trained and untrained cells to perform phagocytosis using attenuated *E. coli* conjugated to a dye which fluoresces in the low-pH environment of a phagosome, allowing us to use fluorescent signal as a readout for phagocytosis efficiency. Both β-Glucan-trained and MDP-trained cells had higher mean fluorescence than untrained cells (indicating more efficient phagocytosis), but MDP-trained cells had 1.1-1.5 times stronger signal than β-Glucan-trained cells 1 hour after stimulation (**Figure 1H**), demonstrating stimulus-specific differences in trained cell function beyond immune gene transcription.

To quantify the strength of trained responses observed across these experiments, we defined a metric called the training capacity, which distills the changes to cells due to memory of a primary stimulus. We calculated training capacity as the log2 fold change of our value of interest (area normalized transcripts per cell for *TNF*, proportion of cells expressing for *IL6* and *CXCL10*, mean fluorescence per cell for phagocytosis) in trained cells over untrained cells from the same donor. A high training capacity indicates a large enhancement due to training, while a training capacity near zero indicates limited training effect (negative values indicate tolerance). Training capacity varied widely across donors (see **Figure 6**). If training were fully nonspecific, the training capacity for β-Glucan and MDP would be the same in the same donor. However, we found that in most donors, the training capacity was higher for one stimulus than the other, with MDP-trained cells consistently showing a higher training capacity (**Figure 1I**). These differences in training capacity for different stimuli eliminate the possibility that there is a single, discrete trained state induced by all primary stimuli.

### The memory state of trained cells is stimulus specific

Once we determined that cells trained with different stimuli did not respond identically to secondary stimulation, we wanted to distinguish between possible explanations for this difference: do different training stimuli simply induce the same memory state to different magnitudes, or do they generate different states entirely?

When testing our trained cells against a bacterial secondary stimulation (LPS, or attenuated *E. coli*), MDP training showed a greater magnitude of change to untrained cells than β-Glucan training in virtually all cases. Thus, one possibility is that there can be degrees of training (and MDP simply induces a higher degree of training than β-Glucan), but the nature of the trained state is the same across both stimuli. If that were the case, then MDP-trained cells <u>would show greater training capacity across every possible phenotypic dimension</u> than β-Glucan-trained cells. However, if the two stimuli generated memory states that were specific to the primary stimulus, there may be contexts in which β-Glucan-trained cells have a greater training capacity than MDP-trained cells (**Figure 2A)**.

**Figure 2:**
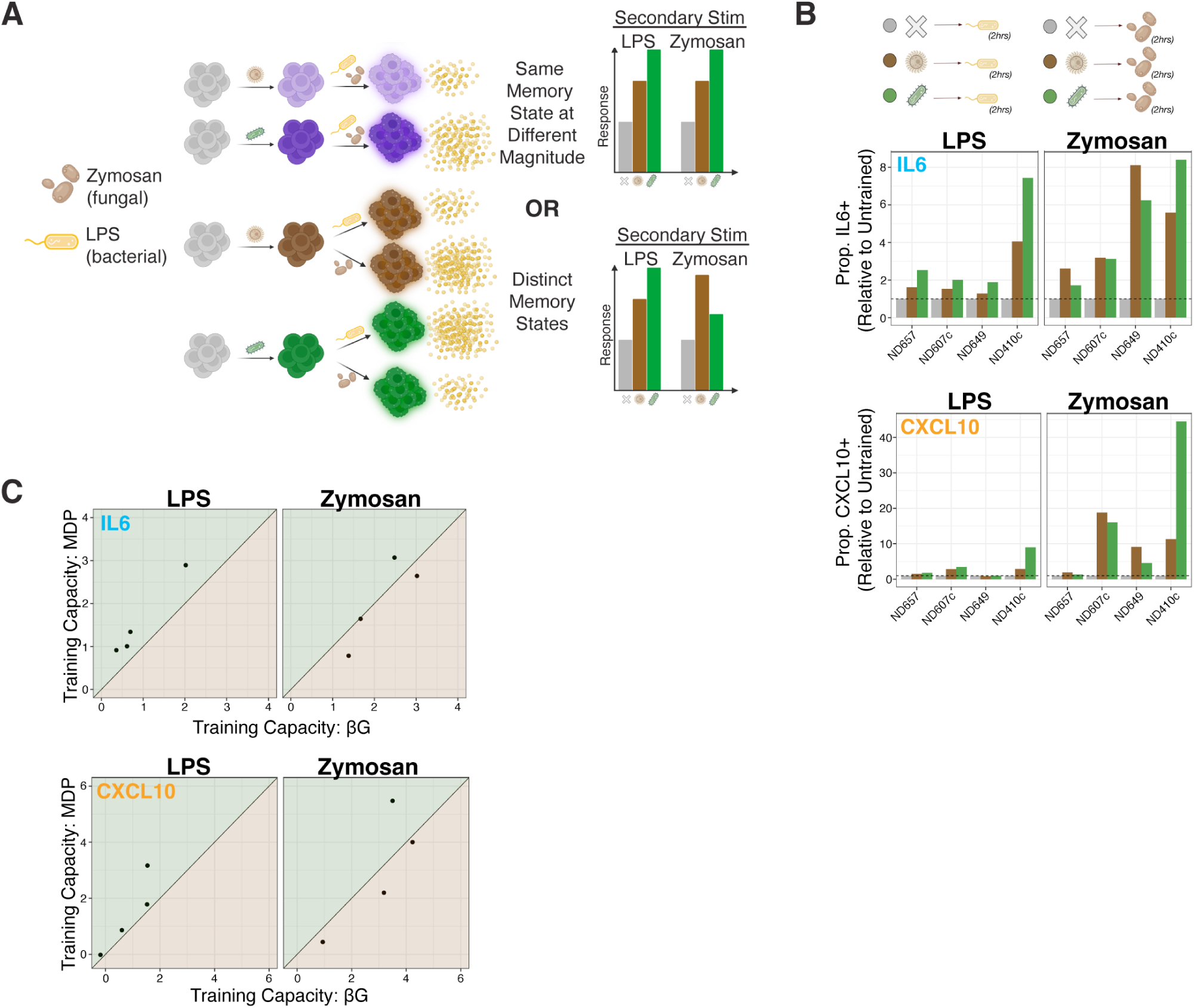
The Memory State of Trained Cells is Stimulus Specific. **(A)** Schematic of experimental question and procedure. If β-Glucan and MDP training exist on a continuum of a universal training state, then one stimulus should always mount a stronger response to secondary stimulation than the other. If memory states are distinct, different stimuli will mount stronger responses to different secondary stimuli. In this experiment, cells were stimulated with either 100ng/mL LPS or 10µg/mL zymosan on day 6. **(B)** Proportion of first responder cells expressing *IL6* or *CXCL10* in untrained (grey), β-Glucan-trained (brown), or MDP-trained (green) population, two hours after stimulation with either LPS or zymosan. Values are normalized to the untrained value for that donor (indicated by dashed line). Approximately 340 single cells were analyzed per condition. **(C)** Summary plots of results from (B), showing training capacity for β-Glucan training versus MDP training. Diagonal *y=x* line denotes where “nonspecific training” would lie, with points above showing stronger training from MDP, and points below showing stronger training from β-Glucan.

**Figure 3:**
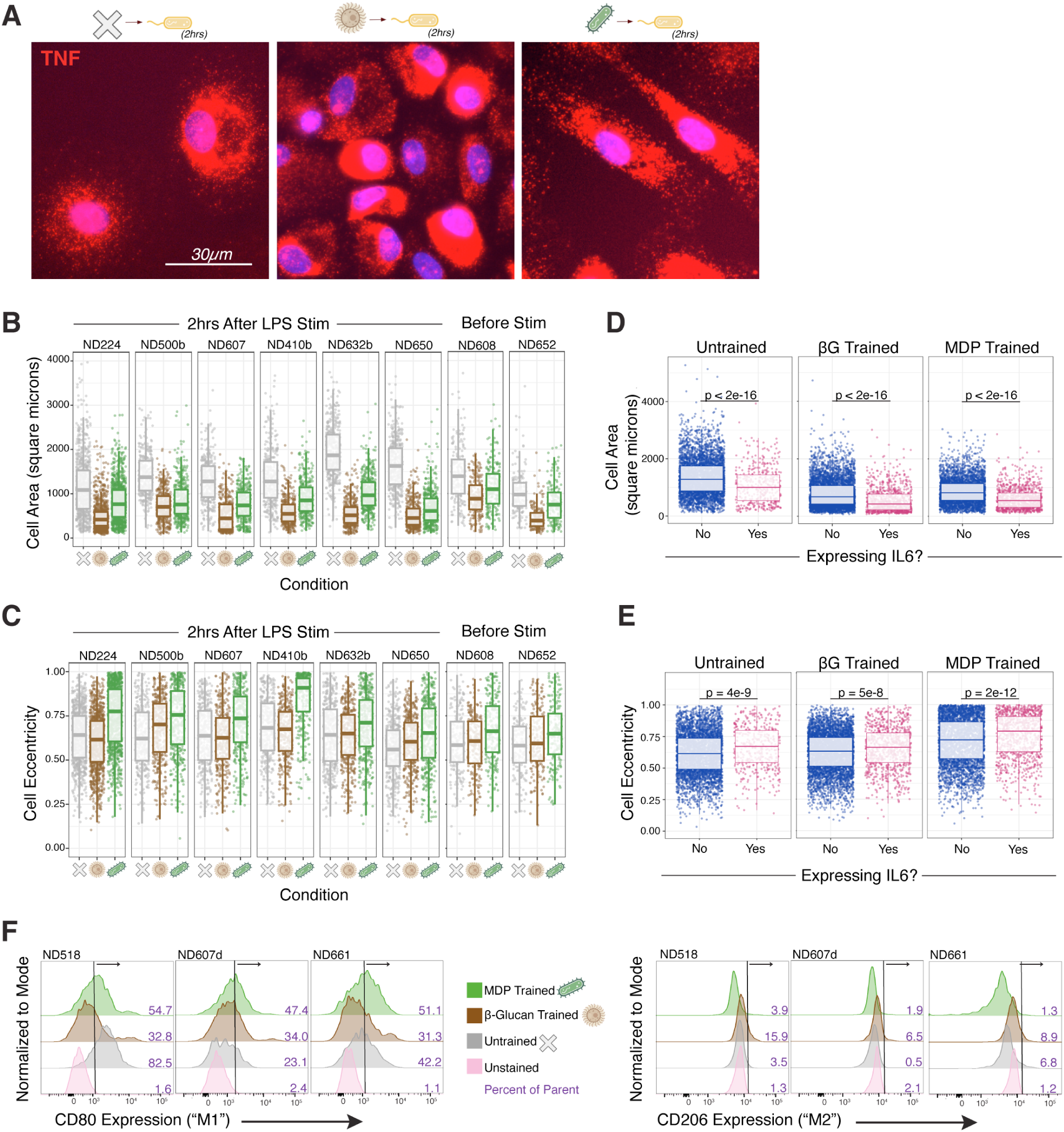
Stimulus-Specific Differences in Trained Cells are Evident Before Restimulation. **(A)** Representative images of single-molecule RNA FISH images of *TNF* expression two hours after stimulation with 100ng/mL LPS, demonstrating distinct morphologies for β-Glucan-trained and MDP-trained cells. **(B)** Cell area (in square microns) for untrained (grey), β-Glucan-trained (brown), and MDP-trained (green) populations before and two hours after stimulation with 100ng/mL LPS. Approximately 415 single cells were analyzed per condition. **(C)** Cell eccentricity for untrained (grey), β-Glucan-trained (brown), and MDP-trained (green) populations before and two hours after stimulation with 100ng/mL LPS. Approximately 415 single cells were analyzed per condition **(D-E)** Differences in cell morphology (area and eccentricity) for cells expressing *IL6* or not expressing *IL6* two hours after stimulation with 100ng/mL LPS, in trained and untrained cells. Data concatenated across 10 total donors. **(F)** Flow cytometry for surface proteins CD80 and CD206, on cells six days after training (but without any secondary stimulation). Histograms show fluorescent intensity normalized to mode for each sample, with unstained control cells in pink. Text denotes percent of gated cells that showed positive signal beyond that of unstained controls.

To test this possibility, we compared the response of β-Glucan-trained and MDP-trained cells to LPS (derived from bacteria, as is MDP) versus zymosan (derived from fungus, as is β-Glucan). In 3 of 4 donors, we found that MDP-trained cells responded more strongly to LPS, while β-Glucan-trained cells responded more strongly to zymosan (**Figure 2B-C, Supplemental Figure 2D**). These results eliminate the possibility that different stimuli are merely eliciting different degrees of a universal activated state and suggest that specificity may allow for heightened responses to a secondary stimulus that is more similar to the inducing memory stimulus, as cells learn to respond in a way more likely to be beneficial based on the information retained about the type of initial stimulation.

### Stimulus-specific differences in trained cells are evident before restimulation

The trained cell phenotype following secondary stimulation differed based on the primary stimulus which was experienced six days prior. We wondered what possible differences may be observable between these populations *before* any secondary stimulation (and were thus associated with the memory state itself).

We first compared cell morphology, which can often predict macrophage state and function, between trained and untrained cells (**Figure 2A**). We immediately noticed that the morphologies of cells trained with different stimuli were qualitatively distinct. β-Glucan-trained cells were smaller (unlike previously reported^13,32^) than untrained cells, while MDP-trained cells were more eccentric than untrained cells. Importantly, these morphologic differences were present before secondary stimulation with LPS, suggesting they were associated with the cell’s memory state as opposed to the response to secondary stimulation (**Figure 2B-C)**. Furthermore, the qualitative differences in morphology between β-Glucan-trained and MDP-trained cells suggest the possibility of specificity in the memory state based on initial stimulus. In both untrained and trained populations, we found that the individual cells that were smaller and more eccentric were more likely to be *IL6* first responders, suggesting that these changes to macrophage morphology due to training may be linked to a predisposition towards inflammatory activation (**Figure 2D-E)**.

Macrophage state is often defined by classification into “M1” polarization (classically activated, often with more eccentric morphology) or “M2” polarization (alternatively activated, often with more rounded morphology)^36,37^. We performed flow cytometry six days after training (but before any restimulation) to quantify expression of cell surface proteins associated with these polarization states. In 2 of 3 donors, a higher proportion of MDP-trained cells expressed CD80 (a marker of M1 state) than untrained cells. In all 3 donors, a higher proportion of β-Glucan-trained cells expressed CD206 (a marker of M2 state) than untrained cells (**Figure 2F, Supplemental Figure 3).**

### Training with host-derived signal is stimulus-specific

IFNγ is a potent, host-derived proinflammatory cytokine known to alter macrophage state at short timescales (24-48hrs, via ‘priming’^24,38,39^). We used it as a training agent to see whether exposure to it, too, was still remembered 6 days after exposure. Like β-Glucan and MDP-trained cells, IFNγ-trained cells also returned to resting state and were no longer producing inflammatory cytokines six days after exposure (**Supplemental Figure 4B**). However, upon secondary stimulation with LPS, IFNγ-trained cells expressed cytokine and chemokine at levels several orders of magnitude higher than untrained cells, demonstrating that the initial stimulus created a memory that persisted over six days. (**Supplemental Figure 4C-E)**. This boost in gene expression was greater than that of β-Glucan-trained and MDP-trained cells.

It was possible that training with IFNγ also generated a distinct memory state within cells, as was the case for β-Glucan and MDP. Because IFNγ signalling is known to disrupt viral replication^40–42^, we tested whether trained cells were less susceptible to infection with virus, with the hypothesis being that IFNγ-trained cells would be the least likely to be infected. We used live replicating HSV-1 virus, with a fluorescent tag on ICP4, a protein involved in viral replication^43^. Results were variable across donors, but we generally found training slightly reduced the proportion of cells infected by HSV-1 (by approximately 6-11%), with IFNγ-trained cells being the least infected in 2 of 3 donors (**Supplemental Figure 4G)**.

### Different stimuli impart distinct memories in chromatin prior to restimulation

Once we had demonstrated that memory “decoding” into phenotype was stimulus-specific, we wanted to connect those phenotypic differences back to the “encoding” of memory into the epigenome. We performed ATAC (assay for transposase-accessible chromatin) sequencing, which measures chromatin accessibility, on cells six days after training with a primary stimulus of β-Glucan, MDP, or IFNγ but before secondary stimulation **(Figure 4A)**. We compared regions of differential accessibility to untrained cells across stimuli to determine whether these memory regions were in the same locations (nonspecific memories) or different locations (specific memories) for cells trained with different stimuli.

**Figure 4:**
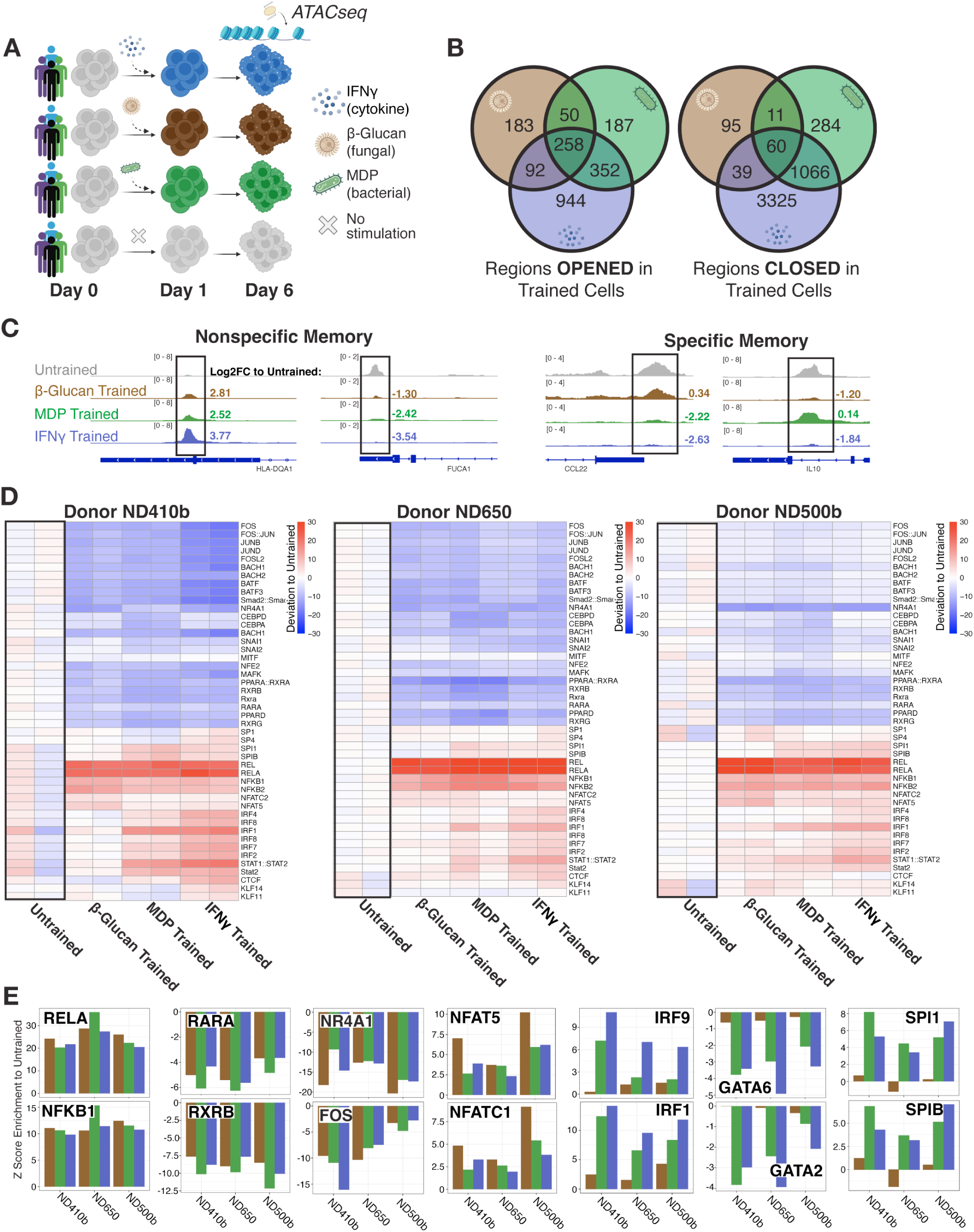
Different Stimuli Impart Distinct Memories in Chromatin Prior to Restimulation. **(A)** Schematic of experimental procedure. Human monocytes were trained with either host cytokine IFNγ, fungal protein β-Glucan, or bacterial mimetic MDP on the first day of culture, before resting for five days. On day 6, cells were harvested for ATAC sequencing to assess chromatin accessibility. **(B)** Venn diagrams of chromatin accessibility changes compared to untrained cells, denoting overlapping regions (nonspecific) as well as unique (specific) regions that change in trained cells in both of the two donors with the strongest training (ND410b and ND650). A log2 fold change cutoff of 1 was used for increasing accessibility, and a cutoff of −1 was used for decreasing accessibility. **(C)** Representative examples of nonspecific (***left***) and specific (***right***) memory regions. At nonspecific memory regions, accessibility changed compared to untrained cells in all three training conditions. At specific memory regions, accessibility changed compared to untrained cells in some, but not all, training conditions. Donor shown is ND410b. **(D)** Heatmap of 46 differential transcription factor motifs for trained and untrained samples from each donor, assessed by *chromVAR*. Heatmap colors denote Z score enrichment for each transcription factor motif in each sample, with untrained cells from that donor set as baseline for comparisons. **(E)** Z score enrichment for transcription factor motif in each sample compared to untrained, for several transcription factors of interest.

We identified both specific and nonspecific regions of memory in trained cells **(Figure 4B-C)**. Two of our three profiled donors (ND410b and ND650) showed ample differences in accessible chromatin in trained cells compared to untrained, while one donor (ND500b) showed more modest changes. For differentially accessible regions found in our two most “trainable” donors, we identified 348 nonspecific memory regions (0.5% of 65,069 total peaks shared between these donors; 258 increasing in accessibility across all three training conditions compared to untrained, 90 decreasing). Of all identified memory regions in β-Glucan-trained cells, 35% were unique to β-Glucan-trained cells (and not differential in either MDP- or IFNγ-trained cells). Of all MDP memory regions, 21% were unique to MDP. Of all IFNγ memory regions, 70% were unique to IFNγ. Memory regions were somewhat enriched in intronic and intergenic sequences, suggesting they may play roles as enhancers (genomic regions which can fine-tune expression of target genes) (**Supplemental Figure 5A)**. This enrichment aligns with prior work demonstrating the importance of enhancer regions in innate immune memory^24,25,44,45^.

Next, we performed a motif analysis looking at variability in accessibility across all peaks in our three donors. In this way, we could determine which transcription factor binding motifs were enriched in regions with differential accessibility in some conditions as compared to untrained **(Figure 4D-E)**. We first looked for transcription factor motifs that were differentially enriched compared to untrained cells across conditions; i.e., general response motifs. We saw a strong positive enrichment for motifs associated with NF-kB across all conditions, fitting with its canonical role as a mediator of the inflammatory response, and as a broad regulator of chromatin remodeling during inflammation^46^. We also saw strong negative enrichment in motifs for transcription factors linked to cell differentiation and immune regulation, including RARA, RXRB, and NR4A1. In 2 of 3 donors, we saw strongly decreased accessibility in regions containing AP-1-associated motifs, including FOS and JUN.

We then asked which transcription factor binding sites (within the above total set of open chromatin regions) were more accessible in one condition than another (specificity regions). β-Glucan-trained cells showed slightly higher enrichment for NFAT motifs, supported by results which show that β-Glucan activates NFAT^47^. Interestingly, MDP- and IFNγ-trained cells uniquely showed differential enrichment in interferon and GATA signaling factor motifs, which were not enriched in β-Glucan trained cells. We also saw much stronger positive enrichment in the macrophage lineage factors SPI1 and SPIB in MDP- and IFNγ-trained cells.

We also performed ATAC sequencing on macrophages six days after training with other primary stimuli (CpG, Poly(I:C), oxLDL, and TGF-β). These cells also showed some hints of specificity in memory, in that their chromatin state was dependent on the stimulus they experienced. OxLDL and TGF-β trained cells showed reduction in NF-kB associated motifs, while cells trained with Poly(I:C) showed strong enrichment for IRF and STAT motifs, but not NF-kB motifs. (Cells trained with CpG showed very limited evidence of memory) (**Supplemental Figure 5B-D)**.

At longer timescales (beyond six days), innate immune memories have been previously shown to be forgotten on a timescale of weeks to months^14,15^. We wondered whether chromatin accessibility similarly showed signs of forgetting at longer times after the initial stimulus, and whether there was stimulus-specificity to any such dynamics. We performed ATAC sequencing on cells from the same donor six days and eleven days after training. Indeed, different stimuli appeared to build more durable memories than others (**Supplemental Figure 6**). Although strongly differential on day six, transcription factor motif enrichment between MDP-trained and untrained cells on day eleven was nearly identical. In contrast, β-Glucan- and IFNγ-trained cells still retained a strong enrichment of inflammatory transcription factor motifs eleven days after training. These findings suggest that training with MDP may be the most likely to be forgotten or rewritten by external environmental cues over time, while training with β-Glucan and IFNγ is more durable. As such, we can identify durability as another facet of innate immune memory which is stimulus-specific, as memory of certain stimuli appears to last longer than others.

Despite differences in chromatin state in trained cells six days after primary stimulation, trained cells returned to a “resting” transcriptional state and were no longer producing proinflammatory cytokines when evaluated (see **Figure 1, Supplemental Figure 1**). As such, these regions of differential chromatin likely represent a memory rather than continued activation stemming from the primary stimulus. Specific differences in transcription factor activity between cells that experienced different stimuli are in line with our phenotypic results, which demonstrate that the specific memories formed during primary stimulation are decoded and lead to different phenotypic behaviors upon a secondary stimulation. Furthermore, these findings suggest a possible mechanism for the encoding of stimulus-specific memories–distinct transcription factors (or cofactors) might be activated by the primary stimulus, binding to distinct genomic regions and leaving behind changes in accessibility that persist for several days after the initial exposure. These results are in line with previous findings that suggest memory formation involves the combination of general stress factors and stimulus-specific response factors^48,49^.

### Any single cell can train

We noted ample variation in levels of cytokine transcription across individual cells in both trained and untrained conditions. As such, we wondered whether every individual cell was capable of remembering the initial stimulation, or only an elite subset of “trainable” cells were able to do so, as has been suggested previously^31^. If only some cells could train, greater cytokine production in the trained population would stem from a small number of cells responding stronger and faster to secondary stimulation, while other cells in the population responded similarly to untrained cells (**Figure 5A**). Conversely, it could be that every cell undergoes training, but that some cells simply do not express cytokine at that moment in time due to other unknown factors^50,51^.

**Figure 5:**
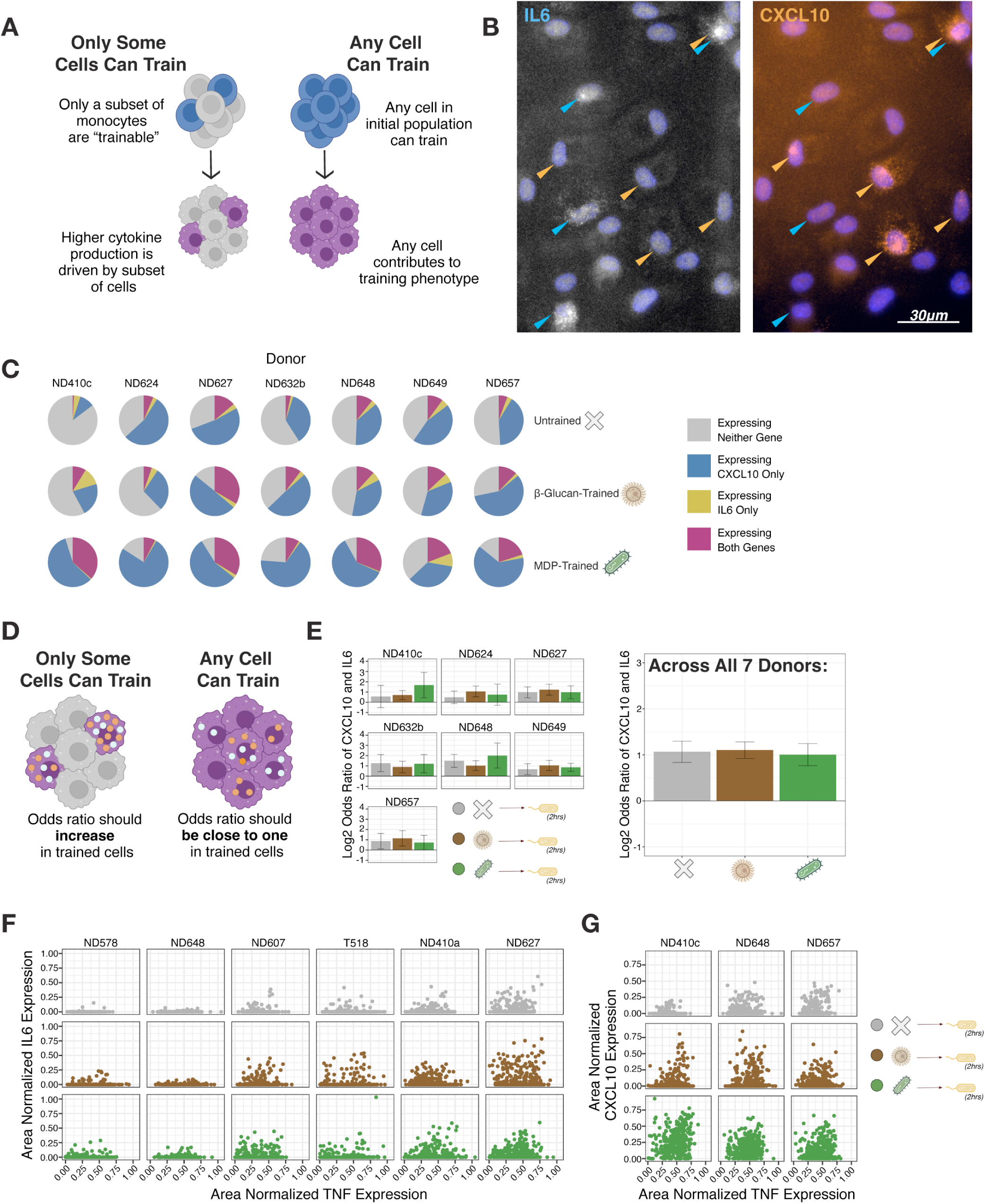
Any Single Cell Can Train. **(A)** Schematic of experimental question. If only a subset of monocytes are trainable, a memory phenotype should only be present in a subset of macrophages six days later. **(B)** Representative RNA FISH image of first responders expressing *IL6* or *CXCL10* two hours after stimulation with LPS, with responding cells indicated by arrows. Donor shown is ND648. **(C)** Proportion of untrained, β-Glucan-trained, and MDP-trained cells expressing *CXCL10, IL6,* neither, or both, two hours after stimulation with LPS. **(D)** Schematic of experimental question. If only some cells can train, there should be an overrepresentation of double positive cells, with a high log2 odds ratio of expression of *IL6* and *CXCL10* in trained cells. If any cell trains, the log2 odds ratio of expression should not increase in trained cells. **(E)** Log2 odds ratio of *CXCL10* and *IL6* expression in untrained (grey), β-Glucan-trained (brown), and MDP-trained (green) cells two hours after stimulation with LPS. Horizontal line denotes a log2 odds ratio of zero. Approximately 300 single cells were analyzed per condition. Left plot shows data concatenated across 7 donors **(F-G)** Area normalized *TNF* versus *IL6,* and *TNF* versus *CXCL10*, expression in untrained (grey), β-Glucan trained (brown), and MDP-trained (green) cells two hours after stimulation with LPS. Each datapoint is one individual cell.

Cytokine expression was extremely heterogeneous across single macrophages in both trained and untrained populations. As such, measuring fold changes in expression of a gene compared to untrained control cells may not be sufficient to identify whether a single cell has trained (especially if population distributions for that gene’s expression between trained and untrained cells are partially overlapping). However, measuring multiple markers of a trained state at once can help to determine whether all cells are capable of training: if different markers of training are present in different cells, that would argue against the existence of an elite subpopulation of cells influenced by the initial training. We therefore returned to our “first responder” cells producing *IL6* or *CXCL10* at two hours after restimulation with LPS (**Figure 5B**).

The proportion of first responders increased in trained populations (**Figure 5C**). If a privileged subpopulation of “trainable” cells existed, we reasoned that these cells would have higher levels of expression of multiple markers of training (e.g. express both *IL6* and *CXCL10* at two hours) compared to untrained cells. At the population level, this pattern would manifest as a higher number of doubly-positive cells than expected by random chance. Quantitatively, the log2 odds ratio of expression for the two genes would be higher in trained cells as compared to untrained cells. However, we instead observed that the log2 odds ratio for expression of *IL6* and *CXCL10* at two hours was around 1.06 (a relatively modest two-fold change), and was nearly identical between trained and untrained cells (**Figure 5D-E**). (First responders for *IL6* were ∼2.5 times more likely to also express *CXCL10* than cells not expressing *IL6*, and first responders for *CXCL10* were ∼1.5 times more likely to also express *IL6* than cells not expressing *CXCL10*). The extremely similar odds ratios in trained and untrained cells indicated there was not an elite subpopulation of trainable cells, suggesting that any single cell in the population was capable of building memories of prior stimulation. We also found that expression of *TNF* vs. *IL6* and *TNF* vs. *CXCL10* was uncorrelated in single cells at these early time points (**Figure 5F-G**), providing further evidence against an elite subset of “trainable” cells driving memory phenotypes in the population.

### Training capacity varies between people and is negatively associated with baseline state

As we investigated differences in memory state across different training stimuli, we also noticed substantial differences in memory across individual donors. We used our summary metric of the training capacity, which quantifies changes to cells due to training, to measure donor variability in our dataset (**Figure 6A**). Donors with a higher training capacity had a larger boost in cytokine expression upon secondary stimulation comparing trained to untrained cells, while those with a lower training capacity showed similar expression levels between trained and untrained cells.

**Figure 6:**
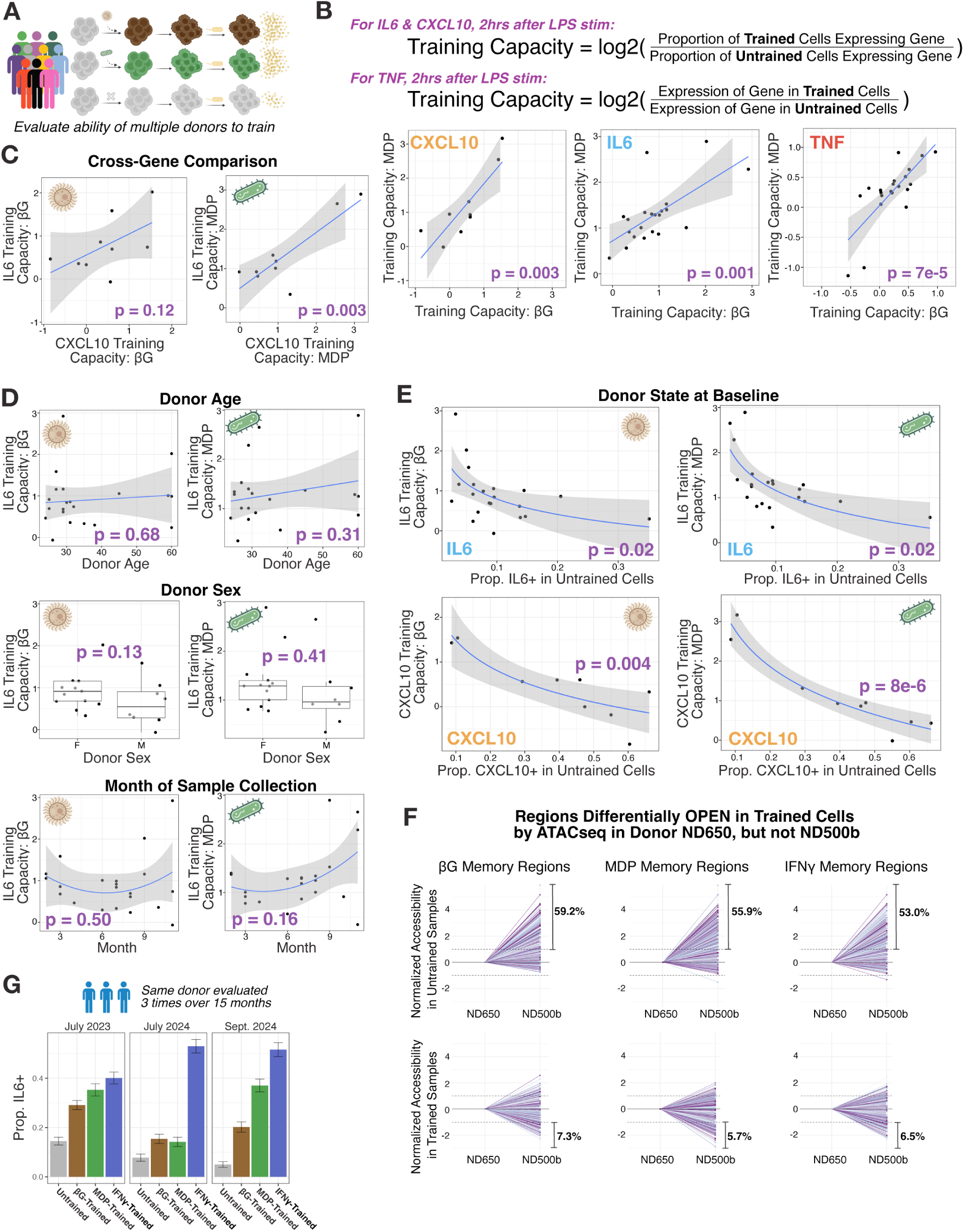
Training Capacity Varies Between People and is Negatively Associated with Baseline State. **(A)** Schematic of question and experimental procedure. Human monocytes from a total of 21 donors were trained with either fungal protein β-Glucan or bacterial mimetic MDP on the first day of culture, before resting for five days. The cytokine response 2 hours after secondary stimulation with LPS was measured by single-molecule RNA FISH. **(B) *Top:*** Equations used to calculate training capacity. ***Bottom:*** Training capacity for β-Glucan versus MDP in the same donor for *IL6, CXCL10,* and *TNF*, and associated p value on linear fit. **(C)** Training capacity for *IL6* versus *CXCL10* in the same donor for β-Glucan and MDP, and associated p value on linear fit. **(D)** Association between training capacity for *IL6* and donor age (***top***, p value on linear fit), donor sex (***middle***, p value on t-test comparison), and month of sample collection (***bottom***, p value on quadratic fit to match seasonal patterns). **(E)** Association between training capacity for *IL6* (***top***) and *CXCL10* (***bottom***) and proportion of cells expressing cytokine in the untrained condition for each donor, and associated p value on a logistic fit. **(F)** ATACseq peaks that were differentially open (log2FC > 1) in trained cells compared to untrained in Donor ND650, but not differential (−1 < log2FC < 1) in Donor ND500b. Each pair of points represents one peak (arbitrary colors), with accessibility normalized by the reads in the ND650 samples. Y-axis shows log2 fold change of the normalized reads in each peak between donors ND500b and ND650. Percentages show the percent of peaks where accessibility is more than twice that of ND650 in ND500b (for untrained cells), or less than half that of ND650 in ND500b (for trained cells). **(G)** Proportion of cells expressing *IL6* in untrained (grey), β-Glucan-trained (brown), MDP-trained (green), or IFNγ-trained (blue) populations, two hours after stimulation with LPS, for the same donor evaluated three times (ND410a, ND410b, ND410c). Error bars show standard error of percentage.

Although β-Glucan and MDP generate different memory states, we found training with either stimulus within the same donor to be strongly correlated (donors that built strong memory of one stimulus also built strong memory of the other) (**Figure 6B**). Within the same donor, training at different loci was also generally correlated (donors with large changes in *IL6* due to training also showed large changes in *CXCL10*) (**Figure 6C**). Training capacity did not seem to depend on donor age or sex, or the time of year of sample collection (**Figure 6D**); however, it is possible that we were not sufficiently powered to observe potential differences.

We did, however, observe a strong association between the untrained (baseline) state of a donor and their training capacity (**Figure 6E**). Macrophages from donors who had a low baseline inflammatory response saw a greater boost in expression from training: the two donors with lowest *IL6* expression in the untrained condition had MDP training capacity of 2.7 and 2.3, while the two donors with highest untrained *IL6* expression had MDP training capacity of 0.56 and 0.92. This trend could suggest that donors with low capacity have preexisting memory of some prior inflammatory experience and are already in an epigenetically “trained” state at time of sample collection. As such, further training would do little to further boost cytokine production (they have reached a “ceiling” of maximal expression).

Our ATAC sequencing data was consistent with this conclusion. For regions that were differentially opened in one donor, but not another, the variation could be due to the trained accessibility being lower in the poorer trainer (indicating less effective training) or due to untrained accessibility being higher in the poorer trainer (indicating a preexisting “trained” chromatin state at baseline). We generally found the latter to be the case (**Figure 6F, Supplemental Figure 7A**). Similar findings have been shown at much larger scale in *in vivo* training with the BCG vaccine in humans^52^, supporting this conclusion.

It’s been suggested that differences in training ability across individuals may be due to genetic differences^53^ or nongenetic differences^52^. To measure the influence of these variables, we profiled cells from the same individual three times over the course of 15 months. We found that this donor’s training capacity varied widely across the three sample dates, and that the baseline untrained value also varied widely over time (**Figure 6G**). We saw similar results for donors profiled twice over the course of several months (**Supplemental Figure 7B**). These findings suggest that transient nongenetic variation plays a larger role than genetic differences in the degree of training in an individual.

## DISCUSSION

Here, we demonstrated stimulus specificity of memory in primary human macrophages days after an initial stimulation. Cells trained with different primary stimuli showed consistent differences in both resting state and in response to secondary challenges. These differences were not simply variations in magnitude of a universal “training” state, but instead reflected distinct memory states specific to each primary stimulus that persisted across several days in culture.

It is known that macrophages dynamically tune their activation state according to a complex network of environmental stimuli, providing context-adjusted responses to stimulation^22,54^. We suggest that specific memories of prior stimulation (not only memory of activation, but of the *type* of activation) also play an important role in macrophage phenotype. Binary categories of “M1” and “M2” macrophages are increasingly viewed as an oversimplification of complex activation states, which can vary widely based on stimulation type and time^55^. Classification schemas that reference specific stimulus memory may improve accuracy and reproducibility of reported macrophage signatures.

What purpose might these specific memories serve? It’s possible that memory provides information to the cell about its prior state, acting as a historical record that allows for adjustment to current responses based on prior experiences. (For example, retaining the information that a specific pathogen type has been experienced recently, allowing for better preparation against the same or similar pathogen). Indeed, it has recently been shown that macrophages stimulated with different ligands at short timescales (∼24 hours) are primed towards a more homogeneous response to different secondary stimulations, although still specific to that initial stimulus^56^. Although governed by entirely different mechanisms to adaptive immunity, trained immunity could provide some of the same benefits to the host of information recording and context-adjusted “learning.”

Although immune cells have evolved specific responses to remember pathogenic stimuli, it has also been demonstrated that cells can remember and learn from experiences with stimuli that the cells have never before experienced in their evolutionary history (such as a targeted inhibitor therapy^57^). In that case, memories were formed via the generic stress factor AP-1 in a “regulation by association” mechanism that allowed cells to dynamically adapt based on prior stressful experiences. It’s possible that similar pathways may play a role in shaping memory specificity in macrophages, as regions of chromatin opened to respond to certain stimuli may be maintained and reinforced, allowing for durable, stimulus specific memories.

We demonstrate specificity of memory in an isolated, *in vitro* setting. It’s possible that specific memories may be overwritten by new environmental cues from other cells in an *in vivo* setting, potentially changing the magnitude or appearance of memory specificity. In a tissue context, these remembered signals also likely occur in combination, making specificities challenging to disentangle. Indeed, we occasionally noted donors who did not follow the stimulus specificity trend we noted on the whole (for example, the two donors who showed significantly higher *IL6* expression in β-Glucan-trained cells than MDP-trained cells). Prior immune memories from a donor’s *in vivo* interactions before cell collection likely influence both the amount of training and the stimulus specificity of that training. Spatial cues may also play a role in memory formation and maintenance, as signal gradients and neighboring cells likely influence what exactly is remembered by each individual cell.

Interestingly, macrophage memory has been shown to be reversible when cells are challenged with a new stimulus^5,58,59^. It remains unknown whether memories of certain stimuli may be more or less mutable. In an *in vitro* setting, we show that memory can persist in primary monocyte-derived macrophages for at least eleven days. Although enhanced cytokine and chemokine production from these memories may be useful in fighting pathogens, accumulation of deleterious memories may influence phenotypes like chronic inflammation. Disentangling stimulus specificities of these memories may unlock vital clues towards generating, or removing, innate immune memories for therapeutic benefit.

## Acknowledgements

The authors would like to thank members of the Raj Lab for scientific discussion and detailed commentary on the manuscript, in particular Pavithran Ravindran, Grant Kinsler, Gianna Busch, and Catherine Triandafillou; Dr. Sunny Shin for expert guidance on macrophage biology; Dr. Shuo Zhang from the Penn Epigenetics Institute, as well as Raj Lab members Vinay Ayyappan and Miles Arnett, for assistance with computational analysis of ATACseq data; summer research students Juan McCook and Nava Graham for scientific discussion and collaborative efforts; the Genomics Sequencing Core at the Wistar Institute, in particular Sonali Majumdar and Sandy Widura, for assistance with sequencing; Professor Nir Drayman for kind gift of HSV1-YCP4 virus; and the William Greenleaf lab for providing custom primer sequences used in ATAC sequencing experiments. Figure cartoons were created using Biorender.com.

The authors thank Emily Cento, Zhilin Chen, Max A. Eldabbas, and Emileigh Maddox of the Human Immunology Core and the Division of Transfusion Medicine and Therapeutic Pathology at the Perelman School of Medicine at the University of Pennsylvania for providing de-identified monocytes that were purified from healthy donor apheresis using StemCell RosetteSep™ kits. The HIC is supported in part by NIH P30 AI045008 and P30 CA016520. HIC RRID: SCR_022380.

A.R. acknowledges support from a center grant from the Mark Foundation for Cancer Research, NIH Director’s Transformative Research Award R01 GM137425, NIH R01 CA238237, NIH R01 CA232256, and NIH 4DN U01 DK127405. A.O. acknowledges support from NSF GRFP DGE-2236662. Z.N. acknowledges support from the Roy and Diana Vagelos Scholars Program in the Molecular Life Sciences, Roy and Diana Vagelos Science Challenge Award, Barry M. Goldwater Scholarship, Fannie and John Hertz Foundation Fellowship, Paul & Daisy Soros Fellowship for New Americans, and Department of Energy Computational Science Graduate Fellowship. J.L. acknowledges support from NIH Medical Scientist Training Program T32GM007170.

## Author Contributions

A.O. and A.R. conceived the project and designed all experiments. Z.N. and A.R. developed image analysis software for all image analysis used in this manuscript, including automated RNA FISH spot detection. J.L assisted A.O. with ATACseq experiments. L.V.E. assisted A.O. with viral infection and flow cytometry experiments. K.S. provided Tn5 enzyme used in ATACseq experiments. A.O. and A.R. prepared all illustrations and wrote the manuscript, with input from all authors.

## Methods

### Key Resources Table

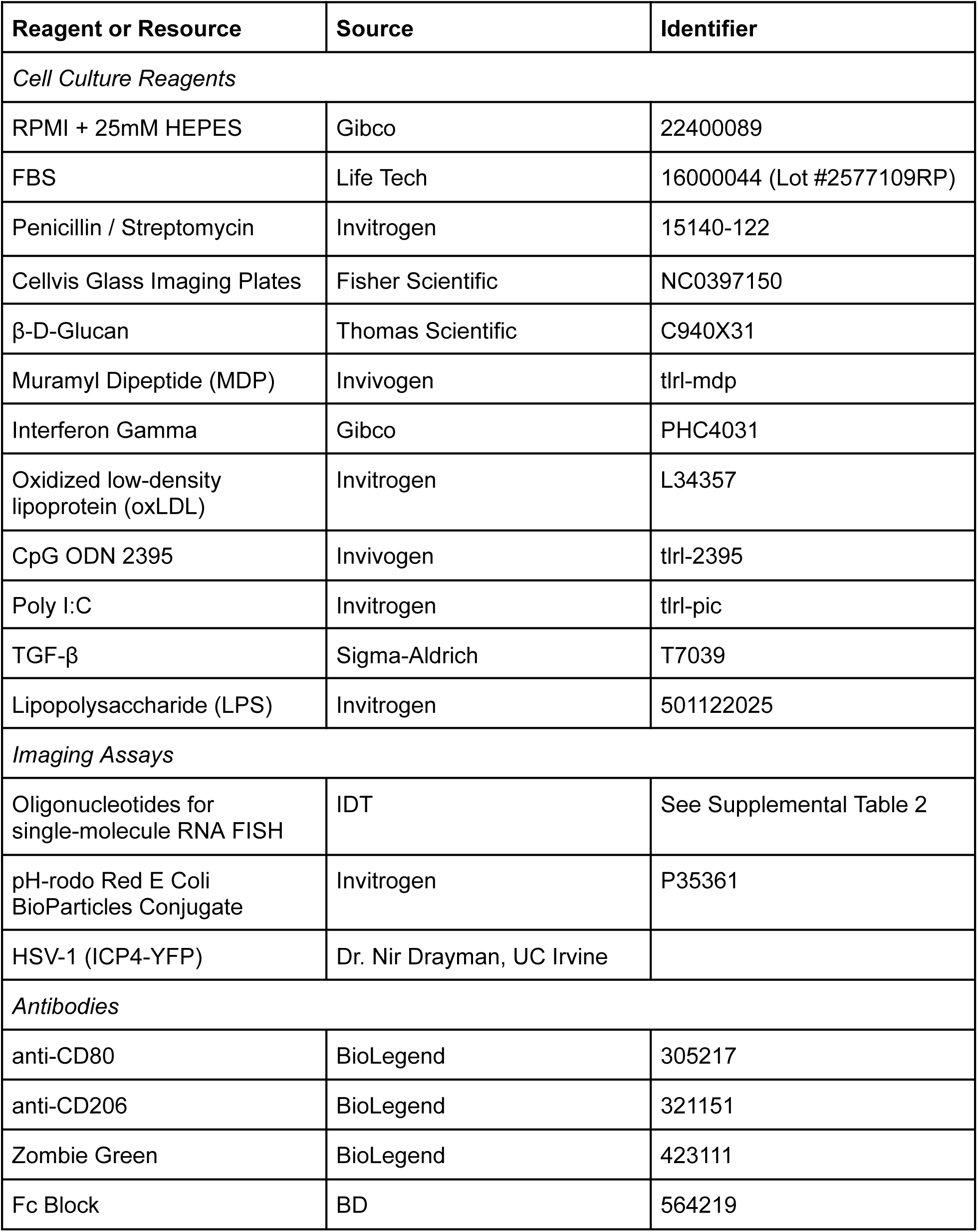
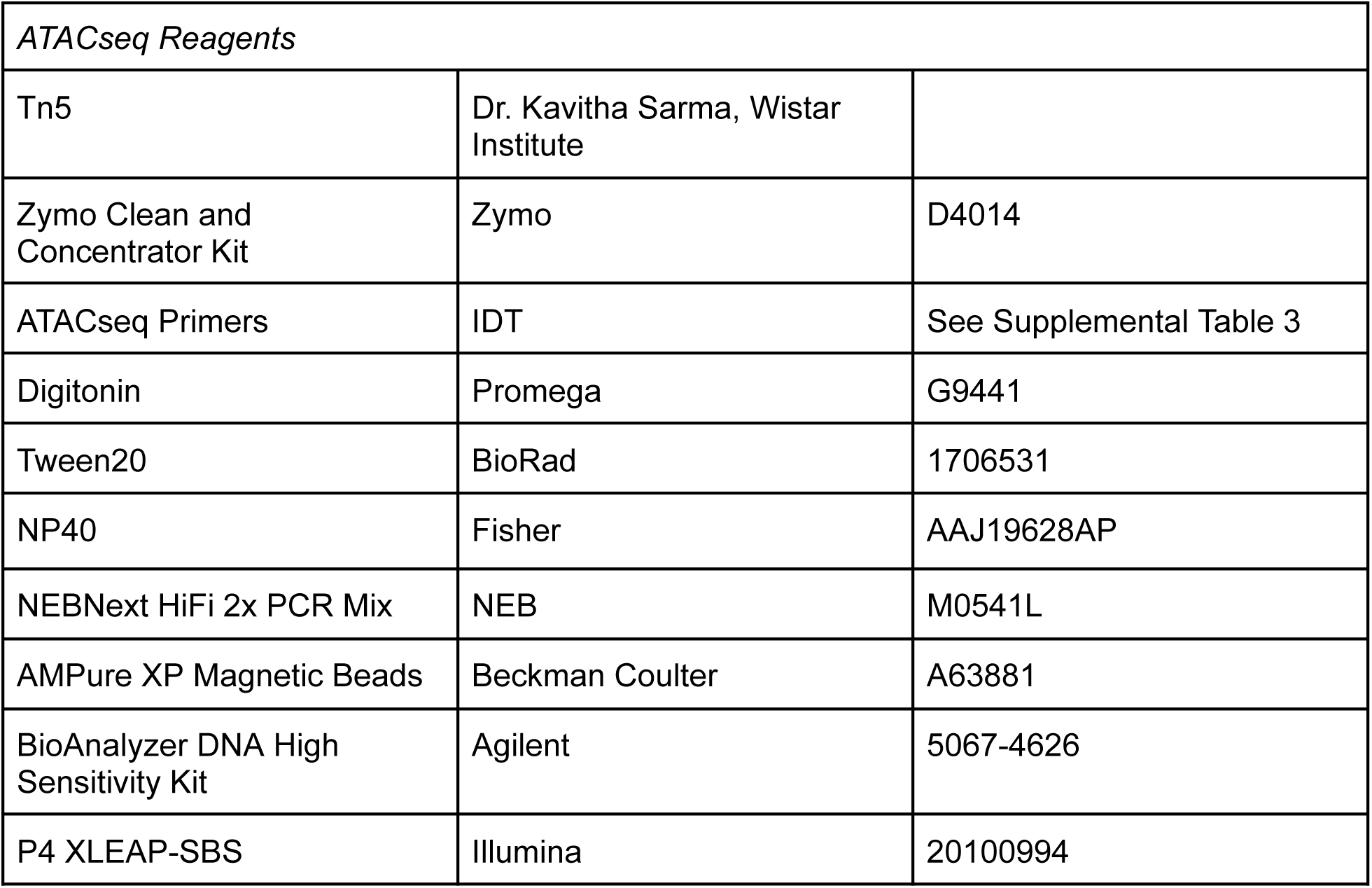

Key Resources Table:
| Reagent or Resource | Source | Identifier |
| --- | --- | --- |
| *Cell Culture Reagents* |  |  |
| RPMI + 25mM HEPES | Gibco | 22400089 |
| FBS | Life Tech | 16000044 (Lot #2577109RP) |
| Penicillin / Streptomycin | Invitrogen | 15140-122 |
| Cellvis Glass Imaging Plates | Fisher Scientific | NC0397150 |
| β-D-Glucan | Thomas Scientific | C940X31 |
| Muramyl Dipeptide (MDP) | Invivogen | tlrl-mdp |
| Interferon Gamma | Gibco | PHC4031 |
| Oxidized low-density lipoprotein (oxLDL) | Invitrogen | L34357 |
| CpG ODN 2395 | Invivogen | tlrl-2395 |
| Poly I:C | Invitrogen | tlrl-pic |
| TGF-β | Sigma-Aldrich | T7039 |
| Lipopolysaccharide (LPS) | Invitrogen | 501122025 |
| *Imaging Assays* |  |  |
| Oligonucleotides for single-molecule RNA FISH | IDT | See Supplemental Table 2 |
| pH-rodo Red E Coli BioParticles Conjugate | Invitrogen | P35361 |
| HSV-1 (ICP4-YFP) | Dr. Nir Drayman, UC Irvine |  |
| *Antibodies* |  |  |
| anti-CD80 | BioLegend | 305217 |
| anti-CD206 | BioLegend | 321151 |
| Zombie Green | BioLegend | 423111 |
| Fc Block | BD | 564219 |

| *ATACseq Reagents* |  |  |
| --- | --- | --- |
| Tn5 | Dr. Kavitha Sarma, Wistar Institute |  |
| Zymo Clean and Concentrator Kit | Zymo | D4014 |
| ATACseq Primers | IDT | See Supplemental Table 3 |
| Digitonin | Promega | G9441 |
| Tween20 | BioRad | 1706531 |
| NP40 | Fisher | AAJ19628AP |
| NEBNext HiFi 2x PCR Mix | NEB | M0541L |
| AMPure XP Magnetic Beads | Beckman Coulter | A63881 |
| BioAnalyzer DNA High Sensitivity Kit | Agilent | 5067-4626 |
| P4 XLEAP-SBS | Illumina | 20100994 |

#### Cell Collection and Culture

De-identified human immune cells were collected from the apheresis product of healthy donors at the Perelman School of Medicine at the University of Pennsylvania by the Human Immunology Core. Monocytes were isolated by negative selection with StemCell RosetteSep kits. Isolated monocytes were cultured in RPMI supplemented with 25mM HEPES, 1% penicillin/streptomycin, and 10% fetal bovine serum, and initially seeded at a density of 800,000 cells per milliliter in Cellvis 24 well glass imaging plates (Fisher NC0397150).

#### Induction of Trained Immunity

We followed an *in vitro* procedure of training commonly used in literature^32^. We introduced training ligands at the following concentrations: β-Glucan at 5 μg/mL, MDP at 1 μg/mL, IFNγ at 50 ng/mL, CpG at 100 ng/mL, Poly I:C at 10 μg/mL, TGF-β at 2 ng/mL, and oxLDL at 10 μg/mL. We chose these doses based on thorough comparison to published literature. Untrained cells received no stimulation during the first 24 hours of culture. After 24 hours, we removed the training ligand by washing all wells (including untrained controls) twice with PBS, then allowed the cells to rest for five additional days, during which time the cells differentiated into monocyte-derived macrophages. On the sixth day of culture, we evaluated trained cells before secondary stimulation, or after stimulation with a readout stimulus. Unless otherwise stated, stimulation on day 6 with lipopolysaccharide (LPS) was at a dose of 100 ng/mL, and stimulation with zymosan was at a dose of 10 µg/mL.

#### Single-Molecule RNA Fluorescence In-Situ Hybridization (FISH)

Oligonucleotides complementary to the transcripts for *TNF*, *CXCL10,* and *IL6* were designed using custom Matlab software (https://github.com/arjunrajlaboratory/ProbeDesign) and purchased from IDT. Due to the short length of these transcripts, we included the 3’ UTR in each target sequence, to generate a total of 24 oligonucleotide probes per gene. To generate fluorescent probes, we first added an amine group to the 3’ end of each oligonucleotide using terminal transferase (TDT), then coupled to either CY3, Alexa 594, or Atto 700 dye.

Single-molecule RNA FISH was performed as described previously^60^. Briefly, cells were fixed in 4% formaldehyde, permeabilized using 70% ethanol, and briefly washed in 5% sodium dodecyl sulfate (SDS) for 1 minute to reduce autofluorescence, before *in situ* hybridization overnight with the probes described in Supplemental Table 2. Cells were then DAPI stained and imaged across 5 Z planes on an inverted Nikon TI-E microscope at 60X magnification.

RNA FISH images were analyzed in custom imaging software NimbusImage (https://github.com/Kitware/UPennContrast). NimbusImage is an open-source, in-browser program that allows for direct visualization and analysis of large imaging datasets. Images were uploaded to NimbusImage, and cell cytoplasm was manually segmented. Single-molecule RNA FISH spots were identified in all Z planes using the deep learning algorithm Piscis^61^, and counted using the “Point Count 3D Projection” tool in NimbusImage. Spot counts of *TNF* were normalized by cell area, and plotted as spots per square micron. Proportions of cells expressing *IL6* and *CXCL10* were calculated using a threshold of 1 spot per cell after manual noise correction.

To calculate statistical significance across proportions of *IL6* and *CXCL10* first responders in trained and untrained cells, we used the following equation: 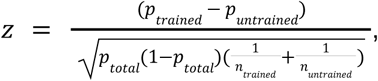 where *p_trained_* is the proportion of first responder cells in the trained population, *p_untrained_* is the proporion of first responders in the untrained population, *p_total_* is the overall sample proportion, and *n_trained_* and *n_untrained_* are the sample sizes. We then used the *pnorm()* function in R to convert the z-score to a p-value, using a two-tailed test for statistical significance.

To calculate statistical significance for *TNF* expression per cell, we calculated the number of spots per square micron in trained and untrained cells, before using the *t.test()* function in R to calculate the p-value for a two-tailed test for statistical significance.

#### Functional Analysis of Phagocytosis and Viral Infection

To evaluate phagocytosis efficacy, trained and untrained cells were stimulated with 100ug/mL of pH-rodo labeled E Coli BioParticles (Invitrogen) for 30 minutes or 1 hour. After stimulation, cells were fixed and DAPI stained, and imaged on an inverted Nikon TI-E microscope at 40X magnification, with fluorescence measured in the Alexa 594 channel. Images were uploaded to NimbusImage, cell cytoplasm was manually segmented, and mean fluorescence per cell was calculated using the “Blob Mean Intensity” tool in NimbusImage.

For viral infection, we used a HSV-1 virus with a YFP fluorescent tag on the ICP4 locus (kind gift of Dr. Nir Drayman, UCI)^43^. Cells were first washed with PBS before infection at a MOI of 2. After 1 hour, the virus was removed and cells remained in culture for 5 additional hours, after which time they were fixed and DAPI stained. Cells were imaged on an inverted Nikon TI-E microscope at 40X magnification, with fluorescence measured in the YFP channel. Images were uploaded to NimbusImage, nuclei segmented using the in-built SegmentAnything tool (https://github.com/facebookresearch/segment-anything) in NimbusImage, and mean fluorescence per nucleus was calculated using the “Blob Mean Intensity” tool in NimbusImage. Infected cells were identified by mean nuclear YFP fluorescence five standard deviations higher than the YFP fluorescence measured in uninfected cells. To calculate statistical significance for difference in proportion of cells infected, we used the following equation:

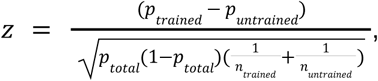

Where *p_trained_* is the proportion of infected cells in the trained population, *p_untrainted_* is the proportion of infected in the untrained population, *p_total_* is the overall sample proportion of infected cells, and *n_trained_* and *n_untranied_* are the sample sizes. We then used the *pnorm()* function in R to convert the z-score to a p-value, using a two-tailed test for statistical significance.

#### Flow Cytometry

Cells were harvested for flow cytometry by first incubating in PBS + 10mM EDTA for 10 minutes on ice, then detaching by vigorous pipetting. Cells were resuspended in Zombie Green Live/Dead stain (1:1000) for 15 minutes, then in Fc block (1:100) for 10 minutes, both at room temperature and protected from light. Antibodies were diluted to a concentration of 1:100 in FACS buffer and incubated with samples for 30 minutes at 4°C, protected from light. Samples were collected on a BD LSR II flow cytometer, and analyzed using FlowJo version 10.8.1.

#### ATAC Sequencing

We performed a modified form of the OMNI-ATAC sequencing protocol described in^62^. We collected 100,000 cells per technical replicate, and lysed in buffer containing 10% NP-40, 10% Tween-20, and 1% Digitonin for 3 minutes at 4°C. Lysed cells were transposed with Tn5 (kind gift of Dr. Kavitha Sarma, Wistar Institute) for 30 minutes at 37°C. Transposed genomic DNA was isolated using a Zymo DNA Clean and Concentrator kit, then amplified using custom primers described in Supplemental Table 3 (originally designed by Greenleaf Lab, Stanford University) for 13 PCR cycles. Libraries were purified using double-sided selection with AMPure XP magnetic beads, and final concentration measured by BioAnalyzer High Sensitivity DNA chips. Libraries were sequenced at depth of approximately 50 million paired-end reads per sample on a NextSeq2000 with a P4 XLEAP 100-cycle kit.

To ensure the highest possible sample quality, we transposed live cells from each donor as they became available. A control sample of MDA-MB-231 cancer cells processed alongside two of these batches showed nearly identical accessibility profiles (**Supplemental Figure 5E**), demonstrating that batch-to-batch variability was minimal.

Reads were first aligned to hg38 using bowtie2. Bam files were cleaned and Tn5 shift was performed using the *bamQC* and *shiftGAlignmentsList* functions from the Bioconductor package ATACseqQC. PCR duplicates were removed using Picard’s *MarkDuplicates* function.

To call peaks, we used the *callpeak* function from MACS2, with an FDR cutoff of 0.05, inputting replicate-merged samples filtered to a fragment size less than 300 base pairs. Once peak regions were identified, we generated one nonredundant peak list shared across all samples and donors, filtered to only include peaks present in at least two samples, and not listed in the ENCODE blacklist regions. We then used GenomicAlignment’s *summarizeOverlaps* function to generate a full counts matrix for all samples (keeping technical replicates separate, and not filtered by fragment size).

To identify differential regions of chromatin, we set a minimum threshold of 50 reads per peak, before normalizing by read depth and averaging the read counts of technical replicates for each peak. We then calculated the log2 fold change between trained and untrained samples, with a cutoff of log2 fold change of 1 for regions opening in trained cells and of −1 for regions closing in trained cells. Regions reported in Figure 3B were differential from untrained cells in both Donor ND410b and Donor ND650. Peak location annotation was identified using *annotatePeak* from the Bioconductor package ChIPseeker^63^.

We used the Bioconductor packages JASPAR2020^64^ and chromVAR^65^ to identify differential transcription factor motif enrichment across samples. The *computeDeviations* function was run on the counts matrix for each donor separately, with *getBackgroundPeaks* and *computeExpectations* set to the untrained samples from that donor to compare trained counts against an appropriate background. The function *deviationScores* was used to calculate z score enrichment for each identified motif.

#### Data and Code Availability

Data and code are accessible at: https://www.dropbox.com/scl/fo/ky20yivgwsnhjl4u0a92z/ADHdeQqMZgoRBWiTPSnoPqI?rlkey=xupdxkeymgnmpd47mzji8viuk&st=j3px6l9l&dl=0

## References

1. Netea, M. G. et al. Trained immunity: A program of innate immune memory in health and disease. Science 352, aaf1098 (2016).

2. Netea, M. G. et al. Defining trained immunity and its role in health and disease. Nat. Rev. Immunol. 20, 375–388 (2020).

3. Divangahi, M. et al. Trained immunity, tolerance, priming and differentiation: distinct immunological processes. Nat. Immunol. 22, 2–6 (2021).

4. Saeed, S. et al. Epigenetic programming of monocyte-to-macrophage differentiation and trained innate immunity. Science 345, 1251086 (2014).

5. Novakovic, B. et al. β-Glucan Reverses the Epigenetic State of LPS-Induced Immunological Tolerance. Cell 167, 1354–1368.e14 (2016).

6. Di Luzio, N. R. & Williams, D. L. Protective effect of glucan against systemic Staphylococcus aureus septicemia in normal and leukemic mice. Infect. Immun. 20, 804–810 (1978).

7. Arts, R. J. W. et al. BCG vaccination protects against experimental viral infection in humans through the induction of cytokines associated with trained immunity. Cell Host Microbe 23, 89–100.e5 (2018).

8. Kaufmann, E. et al. BCG vaccination provides protection against IAV but not SARS-CoV-2. Cell Rep. 38, 110502 (2022).

9. Domínguez-Andrés, J. et al. The Itaconate Pathway Is a Central Regulatory Node Linking Innate Immune Tolerance and Trained Immunity. Cell Metab. 29, 211–220.e5 (2019).

10. Fanucchi, S., Domínguez-Andrés, J., Joosten, L. A. B., Netea, M. G. & Mhlanga, M. M. The Intersection of Epigenetics and Metabolism in Trained Immunity. Immunity 54, 32–43 (2021).

11. Murphy, D. M. et al. Trained immunity is induced in humans after immunization with an adenoviral vector COVID-19 vaccine. J. Clin. Invest. 133, (1 2023).

12. Bekkering, S. et al. Oxidized low-density lipoprotein induces long-term proinflammatory cytokine production and foam cell formation via epigenetic reprogramming of monocytes. Arterioscler. Thromb. Vasc. Biol. 34, 1731–1738 (2014).

13. Ifrim, D. C. et al. Trained immunity or tolerance: opposing functional programs induced in human monocytes after engagement of various pattern recognition receptors. Clin. Vaccine Immunol. 21, 534–545 (2014).

14. Adamson, M. S. et al. RIG-I activation primes and trains innate antiviral immune memory. bioRxiv 2022.10.27.514004 (2022) doi:10.1101/2022.10.27.514004.

15. Kong, L. et al. Single-cell transcriptomic profiles reveal changes associated with BCG-induced trained immunity and protective effects in circulating monocytes. Cell Rep. 37, 110028 (2021).

16. Cheong, J.-G. et al. Epigenetic memory of coronavirus infection in innate immune cells and their progenitors. Cell 186, 3882–3902.e24 (2023).

17. Carlile, S. R. et al. Staphylococcus aureus induced trained immunity in macrophages confers heterologous protection against gram-negative bacterial infection. iScience 27, 111284 (2024).

18. van der Heijden, C. D. C. C. et al. Catecholamines induce trained immunity in monocytes in vitro and in vivo. Circ. Res. 127, 269–283 (2020).

19. Naik, S. et al. Inflammatory memory sensitizes skin epithelial stem cells to tissue damage. Nature 550, 475–480 (2017).

20. Nakayama, Y. et al. Heart failure promotes multimorbidity through innate immune memory. Sci Immunol 9, eade3814 (2024).

21. Sheu, K. M., Guru, A. A. & Hoffmann, A. Quantifying stimulus-response specificity to probe the functional state of macrophages. Cell Syst (2023) doi:10.1016/j.cels.2022.12.012.

22. Luecke, S., Sheu, K. M. & Hoffmann, A. Stimulus-specific responses in innate immunity: Multilayered regulatory circuits. Immunity 54, 1915–1932 (2021).

23. Sheu, K. M., Pimplaskar, A. & Hoffmann, A. Single-cell stimulus-response gene expression trajectories reveal the stimulus specificities of dynamic responses by single macrophages. Mol. Cell 0, (2024).

24. Ostuni, R. et al. Latent enhancers activated by stimulation in differentiated cells. Cell 152, 157–171 (2013).

25. Cheng, Q. J. et al. NF-κB dynamics determine the stimulus specificity of epigenomic reprogramming in macrophages. Science 372, 1349–1353 (2021).

26. Butcher, S. K., O’Carroll, C. E., Wells, C. A. & Carmody, R. J. Toll-Like Receptors Drive Specific Patterns of Tolerance and Training on Restimulation of Macrophages. Front. Immunol. 9, 933 (2018).

27. Kamada, R. et al. Interferon stimulation creates chromatin marks and establishes transcriptional memory. Proc. Natl. Acad. Sci. U. S. A. 115, E9162–E9171 (2018).

28. Tiemeijer, B. M., Heester, S., Sturtewagen, A. Y. W., Smits, A. I. P. M. & Tel, J. Single-cell analysis reveals TLR-induced macrophage heterogeneity and quorum sensing dictate population wide anti-inflammatory feedback in response to LPS. Front. Immunol. 14, 1135223 (2023).

29. Shalek, A. K. et al. Single-cell RNA-seq reveals dynamic paracrine control of cellular variation. Nature 510, 363–369 (2014).

30. Muñoz-Rojas, A. R., Kelsey, I., Pappalardo, J. L., Chen, M. & Miller-Jensen, K. Co-stimulation with opposing macrophage polarization cues leads to orthogonal secretion programs in individual cells. Nat. Commun. 12, 301 (2021).

31. Zhang, B. et al. Single-cell RNA sequencing reveals induction of distinct trained immunity programs in human monocytes. J. Clin. Invest. (2022) doi:10.1172/JCI147719.

32. Bekkering, S. et al. In Vitro Experimental Model of Trained Innate Immunity in Human Primary Monocytes. Clin. Vaccine Immunol. 23, 926–933 (2016).

33. Horneck Johnston, C. J. H., et al. Recognition of yeast β-glucan particles triggers immunometabolic signaling required for trained immunity. iScience 27, 109030 (2024).

34. Kleinnijenhuis, J. et al. Bacille Calmette-Guérin induces NOD2-dependent nonspecific protection from reinfection via epigenetic reprogramming of monocytes. Proc. Natl. Acad. Sci. U. S. A. 109, 17537–17542 (2012).

35. Padovan-Merhar, O. et al. Single mammalian cells compensate for differences in cellular volume and DNA copy number through independent global transcriptional mechanisms. Mol. Cell 58, 339–352 (2015).

36. Rostam, H. M., Reynolds, P. M., Alexander, M. R., Gadegaard, N. & Ghaemmaghami, A. M. Image based Machine Learning for identification of macrophage subsets. Sci. Rep. 7, 3521 (2017).

37. Hourani, T. et al. Label-free macrophage phenotype classification using machine learning methods. Sci. Rep. 13, 5202 (2023).

38. Kang, K. et al. IFN-γ selectively suppresses a subset of TLR4-activated genes and enhancers to potentiate macrophage activation. Nat. Commun. 10, 3320 (2019).

39. Qiao, Y. et al. Synergistic activation of inflammatory cytokine genes by interferon-γ-induced chromatin remodeling and toll-like receptor signaling. Immunity 39, 454–469 (2013).

40. Prestwood, T. R. et al. Gamma interferon (IFN-γ) receptor restricts systemic dengue virus replication and prevents paralysis in IFN-α/β receptor-deficient mice. J. Virol. 86, 12561–12570 (2012).

41. Mikloska, Z. & Cunningham, A. L. Alpha and gamma interferons inhibit herpes simplex virus type 1 infection and spread in epidermal cells after axonal transmission. J. Virol. 75, 11821–11826 (2001).

42. Changotra, H. et al. Type I and type II interferons inhibit the translation of murine norovirus proteins. J. Virol. 83, 5683–5692 (2009).

43. Drayman, N., Patel, P., Vistain, L. & Tay, S. HSV-1 single-cell analysis reveals the activation of anti-viral and developmental programs in distinct sub-populations. Elife 8, (2019).

44. Kaikkonen, M. U. et al. Remodeling of the enhancer landscape during macrophage activation is coupled to enhancer transcription. Mol. Cell 51, 310–325 (2013).

45. Tehrani, S. S. H., Kogan, A., Mikulski, P. & Jansen, L. E. T. Remembering foods and foes: emerging principles of transcriptional memory. Cell Death Differ. (2023) doi:10.1038/s41418-023-01200-6.

46. Feng, A.-C. et al. The transcription factor NF-κB orchestrates nucleosome remodeling during the primary response to Toll-like receptor 4 signaling. Immunity 57, 462–477.e9 (2024).

47. Fanucchi, S. et al. Immune genes are primed for robust transcription by proximal long noncoding RNAs located in nuclear compartments. Nat. Genet. 51, 138–150 (2019).

48. Ghisletti, S. et al. Identification and characterization of enhancers controlling the inflammatory gene expression program in macrophages. Immunity 32, 317–328 (2010).

49. Larsen, S. B. et al. Establishment, maintenance, and recall of inflammatory memory. Cell Stem Cell 28, 1758–1774.e8 (2021).

50. Raj, A. & van Oudenaarden, A. Nature, nurture, or chance: stochastic gene expression and its consequences. Cell 135, 216–226 (2008).

51. Symmons, O. & Raj, A. What’s Luck Got to Do with It: Single Cells, Multiple Fates, and Biological Nondeterminism. Mol. Cell 62, 788–802 (2016).

52. Moorlag, S. J. C. F. M. et al. Multi-omics analysis of innate and adaptive responses to BCG vaccination reveals epigenetic cell states that predict trained immunity. Immunity 57, 171–187.e14 (2024).

53. Messina, N. L., Netea, M. G. & Curtis, N. The impact of human single nucleotide polymorphisms on Bacillus Calmette-Guérin responses. Vaccine 38, 6224–6235 (2020).

54. Gottschalk, R. A. Signaling is the pathway to macrophage function. Trends Immunol. 44, 496–498 (2023).

55. Strizova, Z. et al. M1/M2 macrophages and their overlaps - myth or reality? Clin. Sci. (Lond.) 137, 1067–1093 (2023).

56. Singh, A. et al. Stimulus-response signaling dynamics characterize macrophage polarization states. Cell Syst. 15, 563–577.e6 (2024).

57. Li, J., et al. AP-1 mediates cellular adaptation and memory formation during therapy resistance. bioRxivorg 2024.07.25.604999 (2024).

58. Lavin, Y. et al. Tissue-resident macrophage enhancer landscapes are shaped by the local microenvironment. Cell 159, 1312–1326 (2014).

59. Liu, S. X., Gustafson, H. H., Jackson, D. L., Pun, S. H. & Trapnell, C. Trajectory analysis quantifies transcriptional plasticity during macrophage polarization. Sci. Rep. 10, 12273 (2020).

60. Raj, A., van den Bogaard, P., Rifkin, S. A., van Oudenaarden, A. & Tyagi, S. Imaging individual mRNA molecules using multiple singly labeled probes. Nat. Methods 5, 877–879 (2008).

61. Niu, Z., et al. Piscis: a novel loss estimator of the F1 score enables accurate spot detection in fluorescence microscopy images via deep learning. bioRxiv (2024) doi:10.1101/2024.01.31.578123.

62. Corces, M. R. et al. Omni-ATAC-seq: Improved ATAC-seq protocol. Protoc. Exch. (2017) doi:10.1038/protex.2017.096.

63. Wang, Q., et al. Exploring epigenomic datasets by ChIPseeker. Curr. Protoc. 2, e585 (2022).

64. Fornes, O. et al. JASPAR 2020: update of the open-access database of transcription factor binding profiles. Nucleic Acids Res. 48, D87–D92 (2020).

65. Schep, A. N., Wu, B., Buenrostro, J. D. & Greenleaf, W. J. chromVAR: inferring transcription-factor-associated accessibility from single-cell epigenomic data. Nat. Methods 14, 975–978 (2017).

