## Supplemental Figures and Tables for "Innate Immune Memory is Stimulus Specific"

**Supplemental Information**

**Supplemental Figure 1:**

**(A)** ***Left:*** Representative single-molecule RNA FISH images of *IL6* (top) and *CXCL10* (bottom) expression in trained and untrained cells before secondary stimulation. ***Right:*** Quantified data for representative images, showing area-normalized transcripts per cell. Donor shown for *IL6* is ND608, and for *CXCL10* is ND652.
**(B)** Representative single-molecule RNA FISH image of *TNF* expression two hours after stimulation with 100ng/mL LPS, and associated spot annotations identified using the machine learning algorithm *Piscis.* Donor shown is ND500a.
**(C)** Data shown in Figure 1F-G (proportion of first responder cells expressing IL6 or CXCL10) in untrained (grey), β-Glucan-trained (brown), or MDP-trained (green) populations, two hours after stimulation with LPS), not normalized by the untrained value for each donor. Error bars show standard error of percentage for approximately 400 single cells analyzed per condition. *** p<0.001, ** p<0.01, * p<0.1, NS p>0.1.
**(D)** Data shown in Figure 1E (mean area-normalized TNF transcripts per cell) in untrained (grey), β-Glucan-trained (brown), or MDP-trained (green) populations, two hours after stimulation with LPS, not normalized by the untrained value for each donor. Error bars show standard error of percentage for approximately 400 single cells analyzed per condition. *** p<0.001, ** p<0.01, * p<0.1, NS p>0.1.
**(E)** Mean area-normalized *IL6 (**top**)* or *CXCL10 (**bottom**)* transcripts per cell) in untrained (grey), β-Glucan-trained (brown), or MDP-trained (green) populations, two hours after stimulation with LPS. Values were normalized to the untrained value for that donor (indicated by dashed line).

**Supplemental Figure 2:**

**(A)** Local cell density is not correlated with the proportion of first responders of *IL6* or *CXCL10*, in untrained (grey), β-Glucan-trained (brown), or MDP-trained (green) populations.
**(B)** Proportion of cells expressing *IL6* (***left***) and area-normalized *IL6* transcripts per cell (***right***) in trained and untrained cells, assessed every hour for six hours after stimulation with 100ng/mL LPS. Area under the curve (AUC) of *IL6* expression over time for each population is shown on the plot.
**(C)** Area-normalized *TNF* transcripts per cell in trained and untrained cells, assessed every hour for six hours after stimulation with 100ng/mL LPS. Area under the curve (AUC) of *TNF* expression over time for each population is shown below.
**(D)** Data shown in Figure 2G (proportion of first responder cells expressing IL6 or CXCL10 in untrained (grey), β-Glucan-trained (brown), or MDP-trained (green) populations, two hours after stimulation with LPS or zymosan), not normalized by the untrained value for each donor. Error bars show standard error of percentage for approximately 340 single cells analyzed per condition. *** p<0.001, ** p<0.01, * p<0.1, NS p>0.1.

**Supplemental Figure 3:**

**(A)** Gating strategy for flow cytometry data shown in Figure 2E.

**Supplemental Figure 4:**

**(A)** Schematic of question and experimental procedure. Human monocytes were stimulated with fungal protein β-Glucan, bacterial mimetic MDP, or host-derived cytokine IFNγ on the first day of culture, before resting for five days. The cytokine response to secondary stimulation with heterologous pathogen LPS was measured by single-molecule RNA FISH.
**(B)** ***Left:*** representative images of single-molecule RNA FISH for *TNF* and *IL6* in IFNγ trained cells before any secondary stimulation. ***Right:*** area-normalized transcripts per cell for *TNF* and *IL6* in untrained (grey) and IFNγ-trained (blue) cells before any secondary stimulation. Donor shown is ND608.
**(C-D)** Proportion of first responder cells expressing *IL6* or *CXCL10* in untrained (grey), β-Glucan-trained (brown), MDP-trained (green), or IFNγ-trained (blue) populations, two hours after stimulation with LPS. Values were normalized to the untrained value for that donor (indicated by dashed line). Approximately 450 single cells were analyzed per condition.
**(E)** Summary plots of results from C-D, showing training capacity for each donor in IFNγ versus β-Glucan (***left***) and MDP (***right***) trained cells. Training capacity was calculated by log2 fold change of the difference in proportion of first responders (*IL6, CXCL10*) between trained and untrained cells. Diagonal *y=x* line denotes where "nonspecific training" would lie on these plots, with points above showing stronger training from IFNγ, and points below showing stronger training from β-Glucan or MDP.
**(F)** ***Top:*** Representative single-molecule RNA FISH images of *TNF* expression two hours after stimulation with 100ng/mL LPS, demonstrating distinct morphology in IFNγ trained cells. ***Bottom:*** Cell eccentricity for untrained (grey) and IFNγ-trained (blue) populations two hours after stimulation with 100ng/mL LPS. Approximately 415 single cells were analyzed per condition.
**(G)** ***Left:*** representative image of fluorescent signal from cells six hours after challenge with fluorescently labeled HSV-1. Early stage infection shows diffuse nuclear fluorescent signal, while later stage infection shows punctated signal in the cytoplasm. ***Right:*** Proportion of cells that were infected in untrained (grey), β-Glucan-trained (brown), or MDP-trained (green) populations six hours after viral challenge. Error bars show standard error of percentage for approximately 350 single cells analyzed per condition.

**Supplemental Figure 5:**

**(A)** Region annotation (promoter, UTR, exon, intronic, and intergenic) assessed using ChIPseeker for memory peaks shared between Donors ND410b and ND650.
**(B)** Schematic of experimental procedure. Human monocytes were trained with one of seven possible training agents on the first day of culture, before resting for five days. On day 6, cells were harvested for ATAC sequencing to assess chromatin accessibility.
**(C)** Heatmap of 46 differential transcription factor motifs for trained and untrained samples from each donor, assessed by *chromVAR*. Heatmap colors denote Z score enrichment for each

transcription factor motif in each sample, with untrained cells from that donor set as baseline for comparisons. Donor ND500b can also be found in Figure 4.
**(D)** Z score enrichment for transcription factor motif in each sample compared to untrained, for several transcription factors of interest.
**(E)** ***Left:*** schematic of experimental procedure. MDA-MB-231 cancer cells were grown in culture then frozen down into two separate vials. One vial was processed for ATACseq alongside macrophages from donor ND650, and one vial was processed for ATACseq alongside macrophages from donor ND500b, to assess potential differences across donors that may exist solely due to sample processing. ***Right:*** Normalized reads per peak for frozen controls assessed in each batch. Batches showed strong concordance, with only a very small number of peaks showing differential accessibility across samples.

**Supplemental Figure 6:**

**(A)** Schematic of experimental procedure. Human monocytes were trained with either host cytokine IFNγ, fungal protein β-Glucan, or bacterial mimetic MDP on the first day of culture, before resting. Cells from the same donor were harvested for ATAC sequencing to assess chromatin accessibility on six days and eleven days after training.
**(B)** Z score enrichment for NF-kB family motifs RELA and NFKB1 in each sample compared to untrained.
**(C)** Heatmap of 46 differential transcription factor motifs for trained and untrained samples from each donor, assessed by *chromVAR*. Heatmap colors denote Z score enrichment for each transcription factor motif in each sample, with untrained cells from that sample collection date set as baseline for comparisons. Donor ND410b on Day 6 can also be found in Figure 4.

**Supplemental Figure 7:**

**(A)** ATACseq peaks that were differentially open (log2FC > 1) in trained cells compared to untrained in Donor ND410b, but not differential (-1 < log2FC < 1) in Donor ND500b. Each pair of points represents one peak (arbitrary colors), with accessibility normalized by the reads in the ND410b samples. Y-axis shows log2 fold change of the normalized reads in each peak between donors ND500b and ND410b. Percentages show the percent of peaks where accessibility is more than twice that of ND410b in ND500b (for untrained cells), or less than half that of ND410b in ND500b (for trained cells).
**(G)** Proportion of cells expressing *IL6* in untrained (grey), β-Glucan-trained (brown), MDP-trained (green), or IFNγ-trained (blue) populations, two hours after stimulation with LPS, for the same donor evaluated twice (ND500a vs ND500b, and ND632a vs ND632b). Error bars show standard error of percentage.

**Supplemental Table 1:** De-Identified Donor Information

| **Donor ID** | **Age** | **Sex** | **Date of Collection** |
|---|---|---|---|
| ND224 | 45 | M | 01 February 2024 |
| ND410a | 59 | F | 20 June 2023 |
| ND410b | 60 | F | 11 July 2024 |
| ND410c | 60 | F | 12 September 2024 |
| ND500a | 29 | F | 07 November 2023 |
| ND500b | 30 | F | 18 July 2024 |
| ND518 | 42 | F | 06 November 2024 |
| ND572 | 38 | M | 01 June 2023 |
| ND578 | 32 | M | 30 November 2023 |
| ND580 | 24 | M | 07 February 2024 |
| ND607a | 28 | F | 21 June 2023 |
| ND607b | 28 | F | 05 September 2023 |
| ND607c | 29 | F | 06 August 2024 |
| ND607d | 29 | F | 05 November 2024 |
| ND608 | 27 | F | 19 September 2024 |
| ND616 | 60 | M | 31 August 2023 |
| ND624 | 24 | F | 14 March 2024 |
| ND627 | 31 | M | 19 March 2024 |
| ND632a | 26 | M | 02 November 2023 |
| ND632b | 27 | M | 12 March 2024 |
| ND643 | 29 | F | 08 February 2024 |
| ND648 | 35 | F | 02 July 2024 |
| ND649 | 31 | M | 22 October 2024 |
| ND650 | 25 | F | 16 July 2024 |
| ND651 | 25 | F | 17 October 2024 |

| | | | |
|---|---|---|---|
| ND657 | 25 | F | 08 August 2024 |
| ND658 | 22 | M | 09 October 2024 |
| ND661 | 51 | F | 29 October 2024 |
| T511 | 27 | M | 08 November 2023 |
| T518 | 29 | F | 20 February 2024 |

**Supplemental Table 2**: Oligonucleotide Sequences for RNA FISH Probes

| Gene | Sequence Number | Sequence |
|---|---|---|
| *TNF* | 1 | gggtcagtatgtgagaggaa |
| | 2 | caaagtgcagcaggcagaag |
| | 3 | cggggttcgagaagatgatc |
| | 4 | cttgagggtttgctacaaca |
| | 5 | ctcttgatggcagagaggag |
| | 6 | gatagatgggctcataccag |
| | 7 | caaagtcgagatagtcgggc |
| | 8 | atgatcccaaagtagacctg |
| | 9 | ttgggaaggttggatgttcg |
| | 10 | gtctgaaggagggggtaata |
| | 11 | gtggtcttgttgcttaaagt |
| | 12 | ctgaatcccaggtttcgaag |
| | 13 | ttgaattcttagtggttgcc |
| | 14 | atgtcagggatcaaagctgt |
| | 15 | cattctggccagaaccaaag |
| | 16 | taggtgaggtcttctcaagt |
| | 17 | aaggtccacttgtgtcaatt |

| | | |
|---|---|---|
| | 18 | acatctggagagaggaaggc |
| | 19 | cgtgtctcaaggaagtctgg |
| | 20 | ctacatgggaacagcctatt |
| | 21 | caaaagaaggcacagaggcc |
| | 22 | agtgacagttggtcaccaaa |
| | 23 | gggcgattacagacacaact |
| | 24 | ctttatttctcgccactgaa |
| *IL6* | 1 | gatagagcttctctttcgtt |
| | 2 | agaaggagttcatagctggg |
| | 3 | agaaggcaactggaccgaag |
| | 4 | cggctacatctttggaatct |
| | 5 | cgttctgaagaggtgagtgg |
| | 6 | gtcgaggatgtaccgaattt |
| | 7 | tcacacatgttactcttgtt |
| | 8 | tctttggaaggttcaggttg |
| | 9 | aagcatccatctttttcagc |
| | 10 | ctcctcattgaatccagatt |
| | 11 | tgatgattttcaccaggcaa |
| | 12 | ctggaggtactctaggtata |
| | 13 | tcctcactactctcaaatct |
| | 14 | cttttgtactcatctgcaca |
| | 15 | gggtggttattgcatctaga |
| | 16 | tgagatgagttgtcatgtcc |
| | 17 | aactccttaaagctgcgcag |
| | 18 | catgctacatttgccgaaga |

| | | |
|---|---|---|
| | 19 | acaggtttctgaccagaaga |
| | 20 | aacataagttctgtgcccag |
| | 21 | ctcatacttttagttctcca |
| | 22 | ttcaaactgcatagccactt |
| | 23 | ccaagaaatgatctggctct |
| | 24 | atttgaggtaagcctacact |
| *CXCL10* | 1 | ctgctgtaggctcagaatat |
| | 2 | gattcatggtgctgagactg |
| | 3 | aggcagcaaatcagaatggc |
| | 4 | gaatgccacttagagtcaga |
| | 5 | ccttctttttcattgtagca |
| | 6 | ggattcagacatctcttctc |
| | 7 | aattcttgatggccttcgat |
| | 8 | ttagacctttccttgctaac |
| | 9 | cctctggttttaaggagatc |
| | 10 | tggaagcactgcatcgattt |
| | 11 | gcttgacatatactccatgt |
| | 12 | cacctttttagtgtaactgca |
| | 13 | ctgatttggtgaccatcatt |
| | 14 | acattaaccttcctacagga |
| | 15 | tgccagggtagagttattac |
| | 16 | acctcagtagagcttacatt |
| | 17 | gggtcagaacatccactaag |
| | 18 | agatgggaaaggtgagggaa |
| | 19 | aagattccttagtacccttg |

| | | |
|---|---|---|
| | 20 | ttctgataaaccccaaagca |
| | 21 | cagtggaagtccatgaagta |
| | 22 | gccactgaaagaatttgggc |
| | 23 | acttctactttgtacagtct |
| | 24 | catgttattccatgtacact |

**Supplemental Table 3**: Custom Primers for ATAC Sequencing

| Name | Index | Full Sequence |
|---|---|---|
| N7001 | TAAGGCGA | CAAGCAGAAGACGGCATACGAGATTCGCCTTAGTCTCGTGGGC TCGGAGATGTG |
| N7002 | CGTACTAG | CAAGCAGAAGACGGCATACGAGATCTAGTACGGTCTCGTGGGC TCGGAGATGTG |
| N7003 | AGGCAGAA | CAAGCAGAAGACGGCATACGAGATTTCTGCCTGTCTCGTGGGC TCGGAGATGTG |
| N7004 | TCCTGAGC | CAAGCAGAAGACGGCATACGAGATGCTCAGGAGTCTCGTGGGC TCGGAGATGTG |
| N7005 | GGACTCCT | CAAGCAGAAGACGGCATACGAGATAGGAGTCCGTCTCGTGGGC TCGGAGATGTG |
| N7006 | TAGGCATG | CAAGCAGAAGACGGCATACGAGATCATGCCTAGTCTCGTGGGC TCGGAGATGTG |
| N7007 | CTCTCTAC | CAAGCAGAAGACGGCATACGAGATGTAGAGAGGTCTCGTGGGC TCGGAGATGTG |
| N7008 | CAGAGAGG | CAAGCAGAAGACGGCATACGAGATCCTCTCTGGTCTCGTGGGC TCGGAGATGTG |
| N7009 | GCTACGCT | CAAGCAGAAGACGGCATACGAGATAGCGTAGCGTCTCGTGGGC TCGGAGATGTG |
| N7010 | CGAGGCTG | CAAGCAGAAGACGGCATACGAGATCAGCCTCGGTCTCGTGGGC TCGGAGATGTG |
| N7011 | AAGAGGCA | CAAGCAGAAGACGGCATACGAGATTGCCTCTTGTCTCGTGGGC TCGGAGATGTG |
| N7012 | GTAGAGGA | CAAGCAGAAGACGGCATACGAGATTCCTCTACGTCTCGTGGGC TCGGAGATGTG |

| Name | Index | Full Sequence |
|---|---|---|
| N5001 | TAGATCGC | AATGATACGGCGACCACCGAGATCTACACTAGATCGCTCGTCGG CAGCGTCAGATGTGTAT |
| N5002 | CTCTCTAT | AATGATACGGCGACCACCGAGATCTACACCTCTCTATTCGTCGGC |

| | | |
|---|---|---|
| | | AGCGTCAGATGTGTAT |
| N5003 | TATCCTCT | AATGATACGGCGACCACCGAGATCTACACTATCCTCTTCGTCGGC AGCGTCAGATGTGTAT |
| N5004 | AGAGTAGA | AATGATACGGCGACCACCGAGATCTACACAGAGTAGATCGTCGG CAGCGTCAGATGTGTAT |
| N5005 | GTAAGGAG | AATGATACGGCGACCACCGAGATCTACACGTAAGGAGTCGTCGG CAGCGTCAGATGTGTAT |
| N5006 | ACTGCATA | AATGATACGGCGACCACCGAGATCTACACACTGCATATCGTCGG CAGCGTCAGATGTGTAT |
| N5007 | AAGGAGTA | AATGATACGGCGACCACCGAGATCTACACAAGGAGTATCGTCGG CAGCGTCAGATGTGTAT |
| N5008 | CTAAGCCT | AATGATACGGCGACCACCGAGATCTACACCTAAGCCTTCGTCGG CAGCGTCAGATGTGTAT |
| N5009 | TGGAAATC | AATGATACGGCGACCACCGAGATCTACACTGGAAATCTCGTCGG CAGCGTCAGATGTGTAT |
| N5010 | AACATGAT | AATGATACGGCGACCACCGAGATCTACACAACATGATTCGTCGG CAGCGTCAGATGTGTAT |
| N5011 | TGATGAAA | AATGATACGGCGACCACCGAGATCTACACTGATGAAATCGTCGG CAGCGTCAGATGTGTAT |
| N5012 | GTCGGACT | AATGATACGGCGACCACCGAGATCTACACGTCGGACTTCGTCGG CAGCGTCAGATGTGTAT |

### Supplemental Figure 1

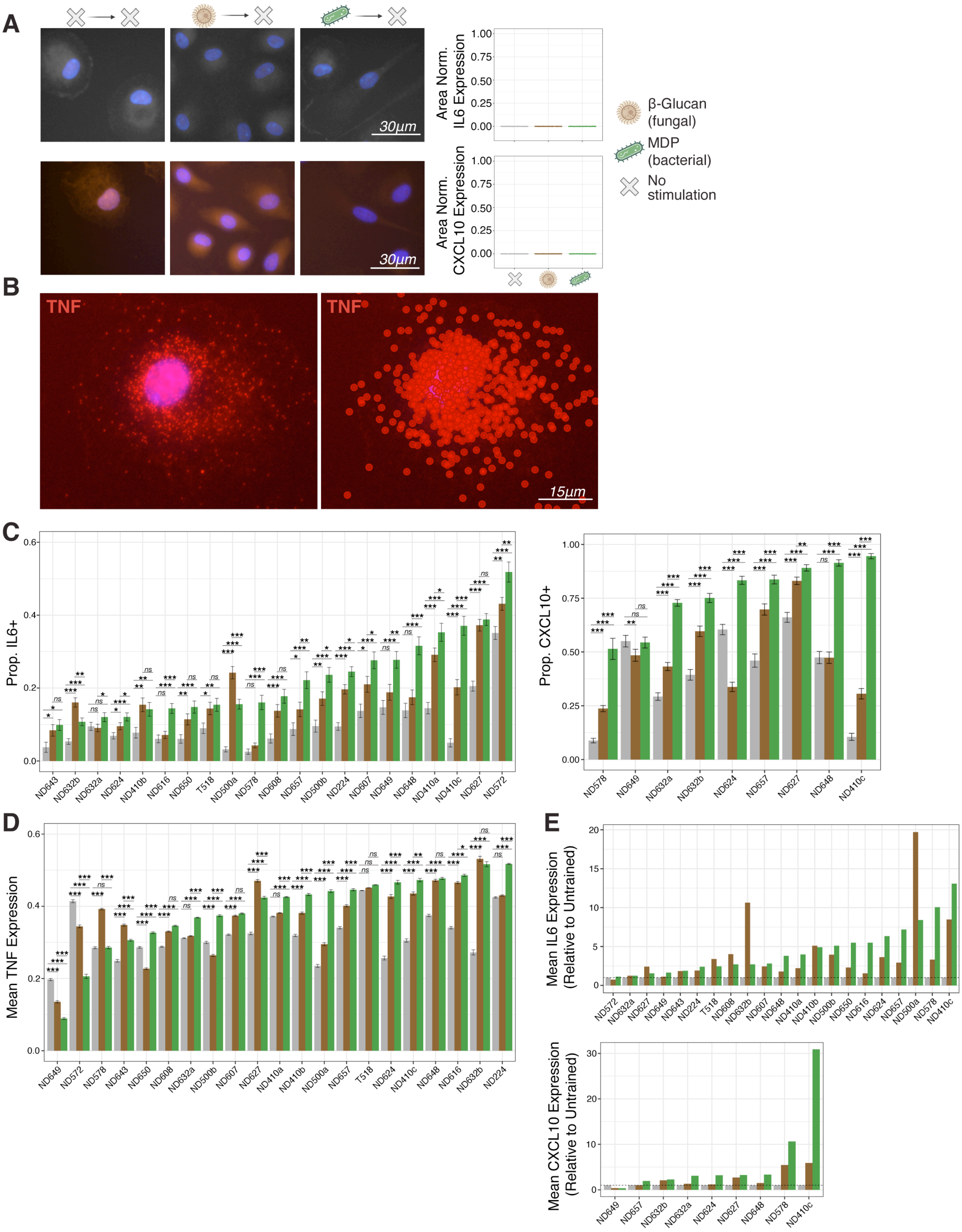

### Supplemental Figure 2

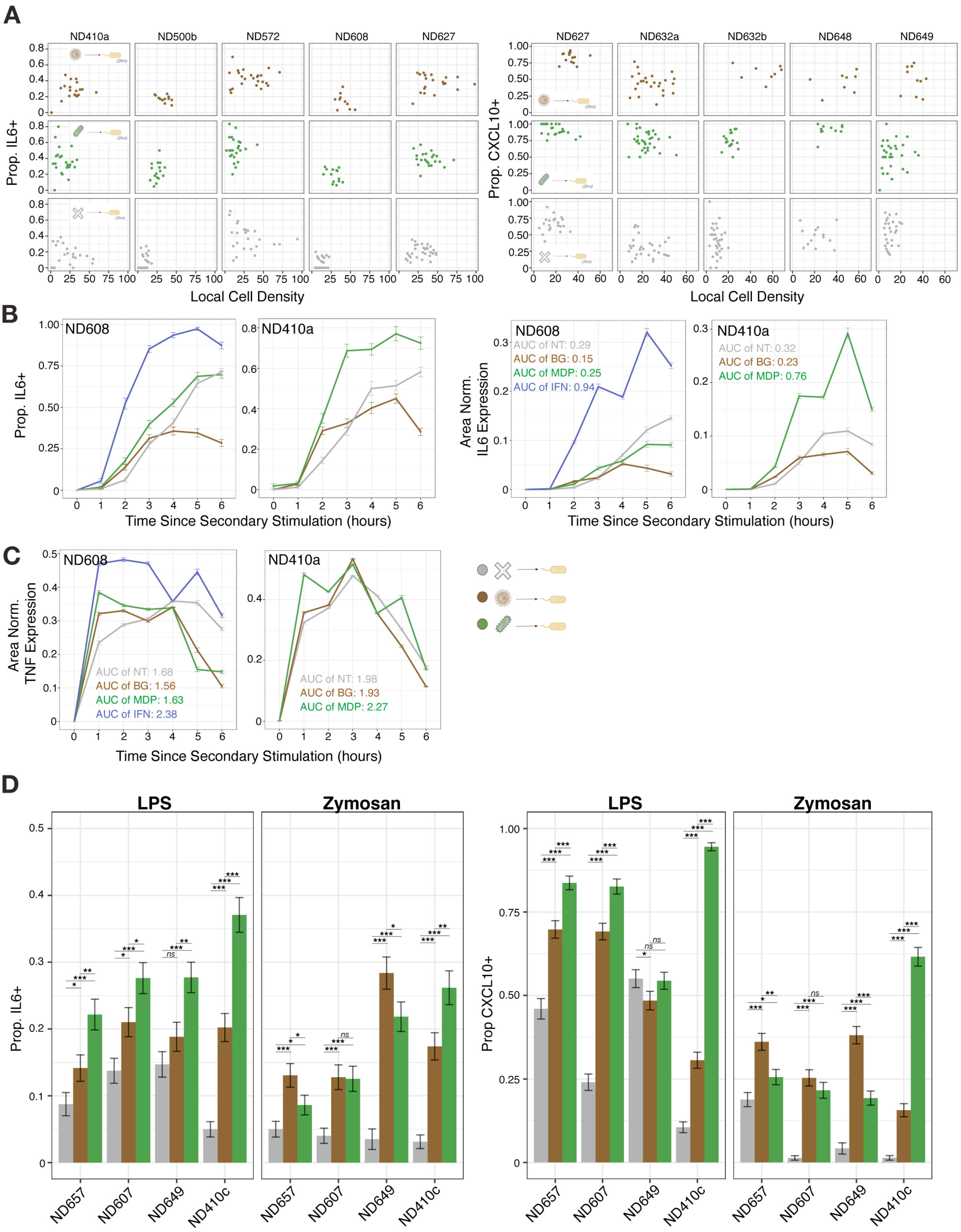

#### Supplemental Figure 3

**A**

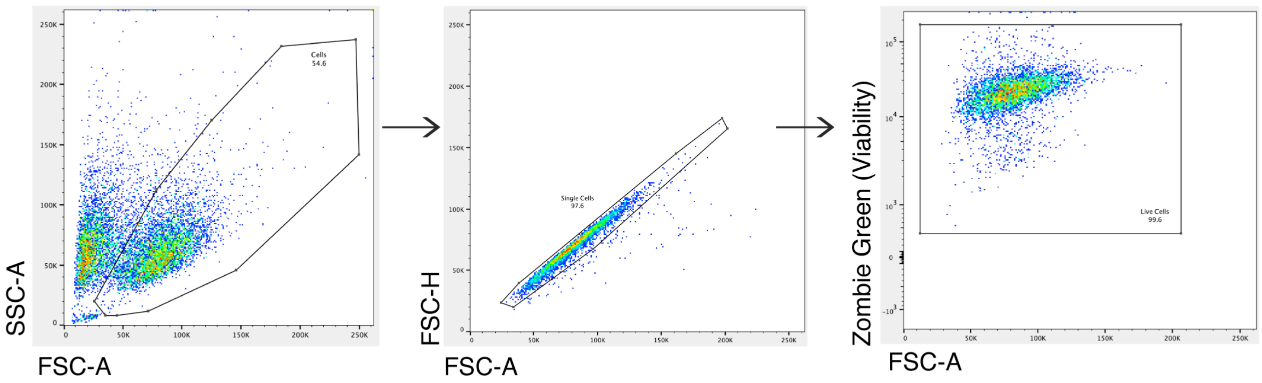

### Supplemental Figure 4

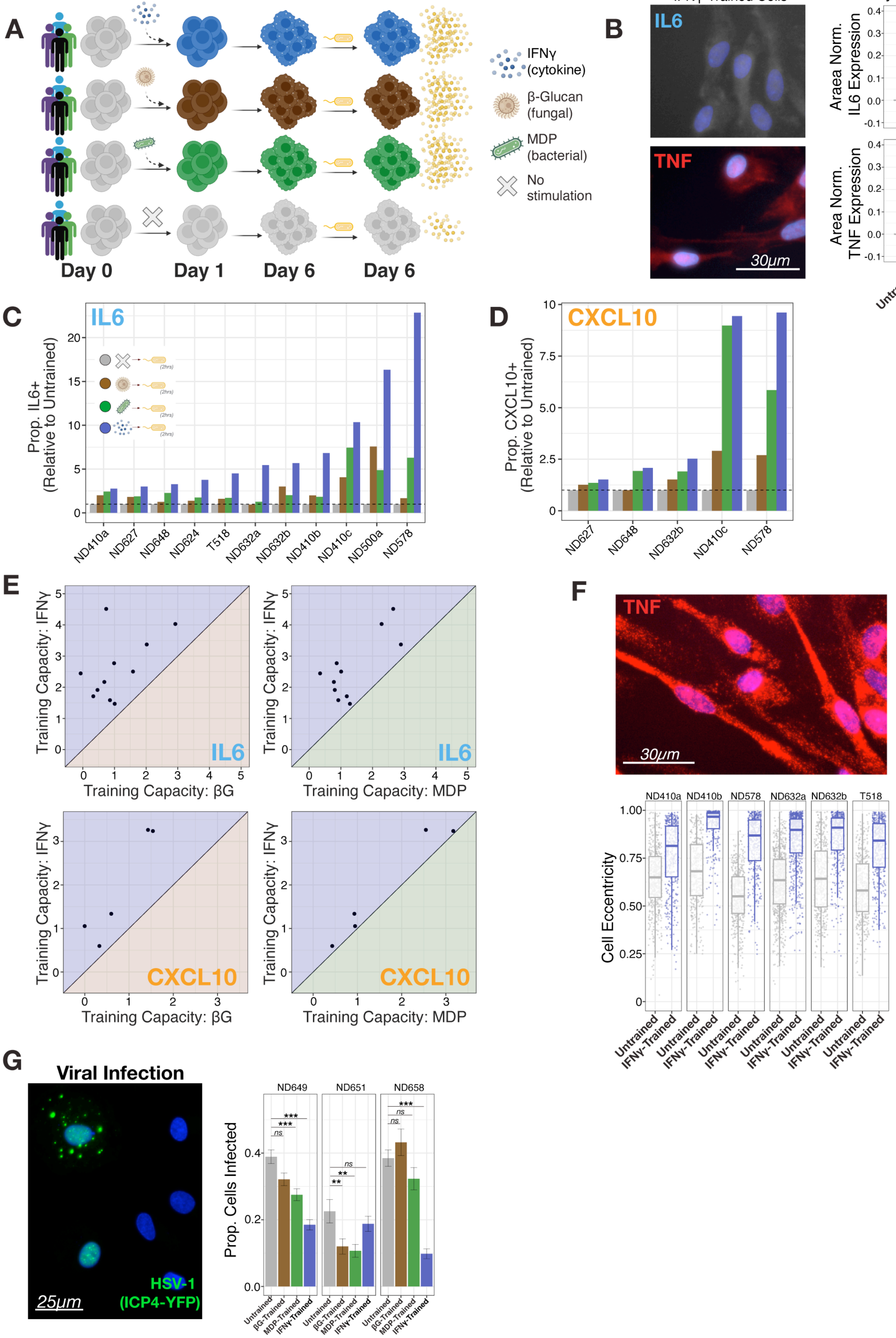

### Supplemental Figure 5

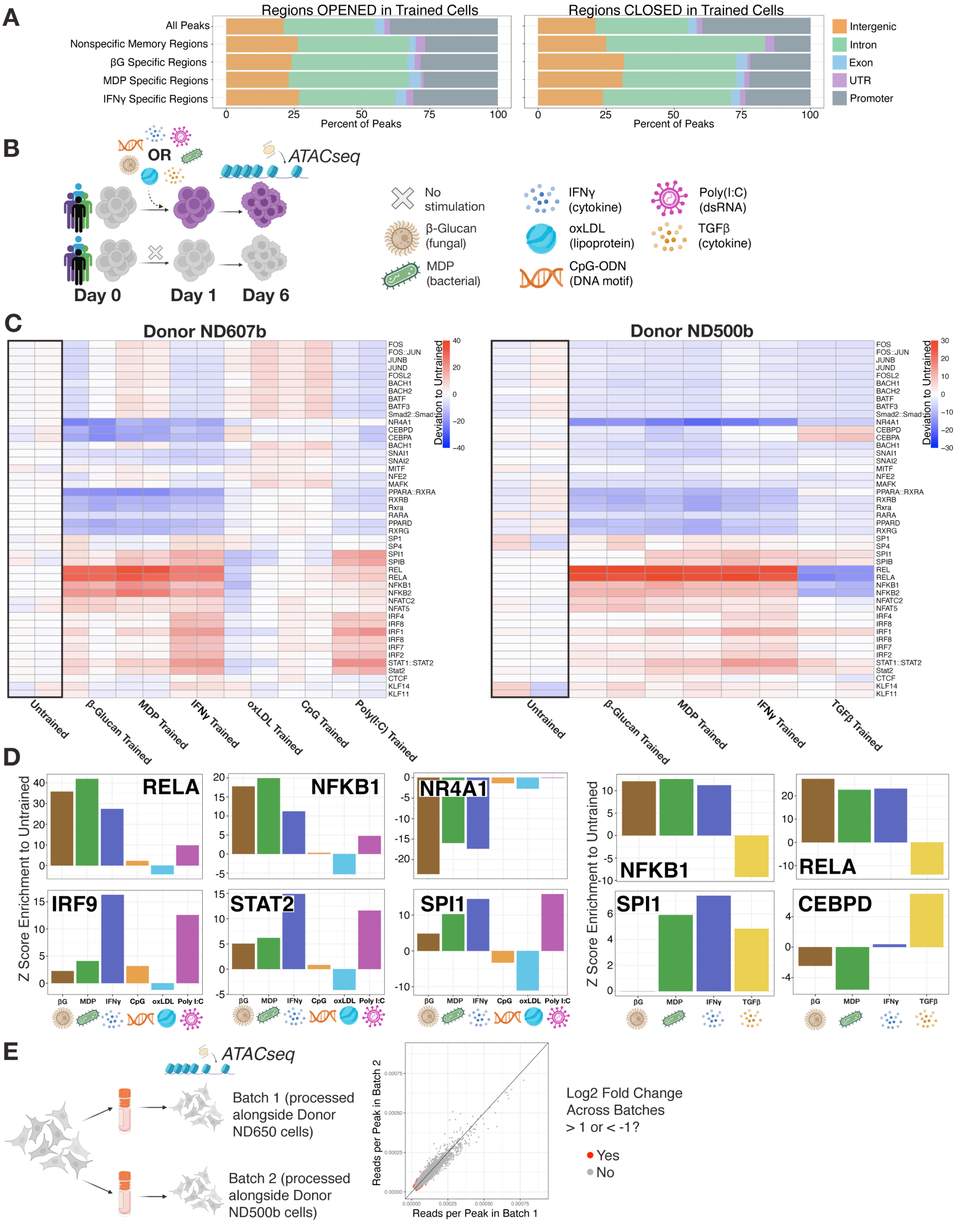

### Supplemental Figure 6

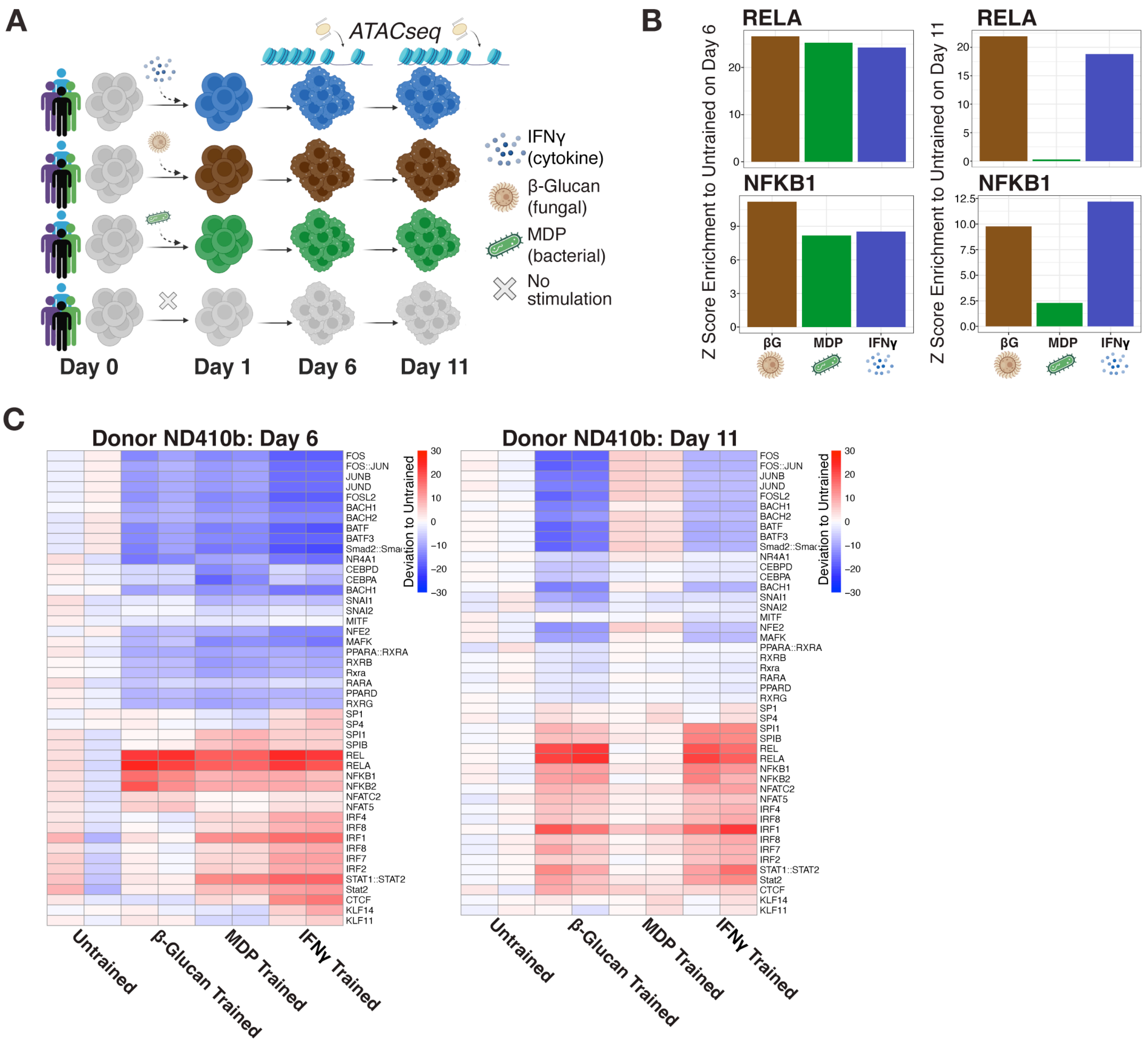

### Supplemental Figure 7

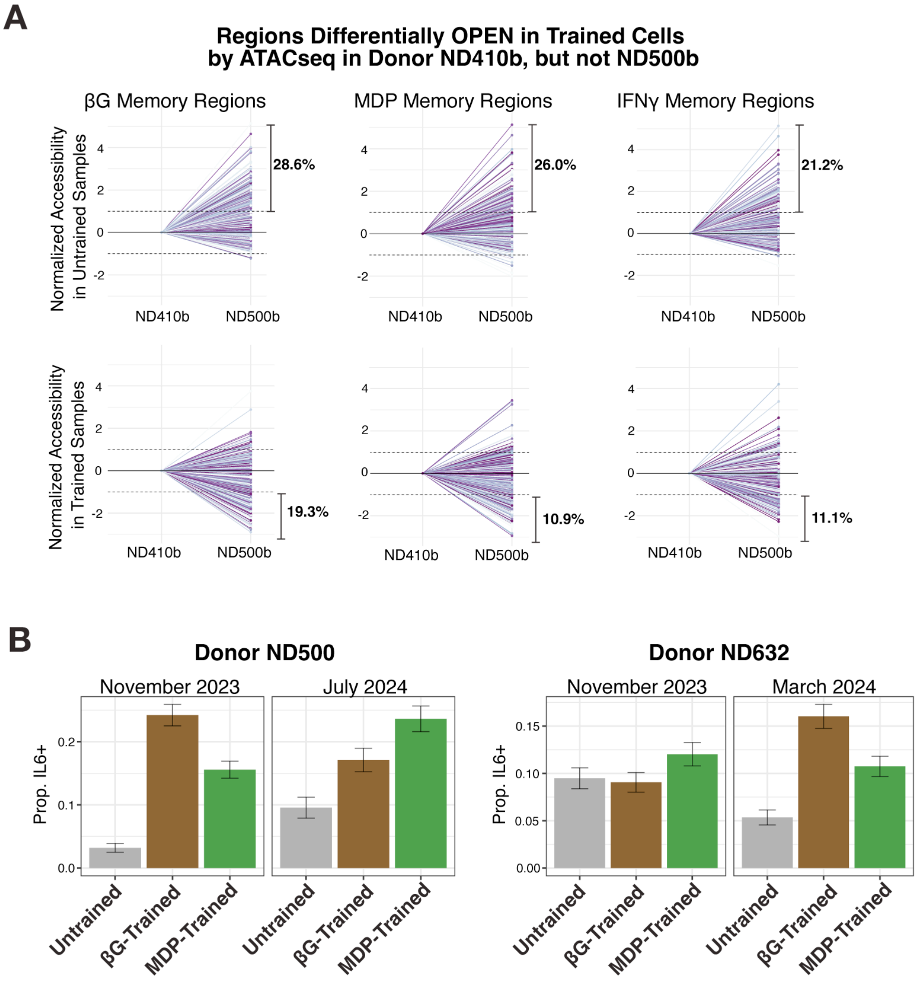
